# Fragment Screening by Weak Affinity Chromatography Identifies Ligands for *Neisseria gonorrhoeae* Undecaprenyl Diphosphate Synthase

**DOI:** 10.1101/2025.05.21.654962

**Authors:** André da Silva Santiago, Bruno Sérgio do Amaral, Edgar Llontopp, Priscila Zonzini Ramos, Lucas Rodrigo Souza, Micael Rodrigues Cunha, Erika Piccirillo, Vitor M. Almeida, Ronaldo Aloise Pilli, Gabriela Barreiro, Katlin B. Massirer, Rafael M. Couñago

**Author notes:** These authors contributed equally to this work (André da Silva Santiago, Bruno Sérgio do Amaral).

## Abstract

The rapid rise of antimicrobial resistance in *Neisseria gonorrhoeae* underscores the urgent need for new therapeutic strategies. Undecaprenyl diphosphate synthase (UPPS), an essential enzyme in bacterial cell wall biosynthesis, has emerged as a promising antimicrobial target. However, no structural information has been available for the gonococcal enzyme (*Ng*UPPS), and known ligand scaffolds remain limited, hindering structure-based drug discovery efforts. Here, we present a combined weak affinity chromatography–mass spectrometry (WAC-MS) and crystallographic pipeline to identify and structurally characterize fragment binders of *Ng*UPPS. WAC-MS screening of a diverse fragment library identified 23 putative binders, 16 of which were successfully co-crystallized with *Ng*UPPS, primarily via soaking into preformed apo crystals. The resulting high-resolution structures mapped fragment binding across four distinct regions of the enzyme’s hydrophobic tunnel, encompassing the pyrophosphate-binding pockets and the distal product-binding site. Additionally, we determined the co-crystal structure of *Ng*UPPS with **BPH-1100**, a known phthalic acid–based UPPS inhibitor, revealing that the enzyme adopts an open, ligand-competent conformation without significant rearrangement. Our results validate *Ng*UPPS as a structurally tractable antibacterial target and provide a strong foundation for fragment-based optimization and the future development of novel inhibitors to combat drug-resistant *N. gonorrhoeae*.

## Introduction

Neisseria-related diseases are caused by two strictly human pathogens: *Neisseria gonorrhoeae* and *Neisseria meningitidis*, the causative agents of gonorrhea and meningitis, respectively. *N. gonorrhoeae*, in particular, accounts for one of the most prevalent sexually transmitted infections worldwide, with a particularly high burden in low- and middle-income countries. Although gonorrhea is associated with low mortality, it can lead to serious reproductive health complications, especially in women, due to its frequently asymptomatic course of infection.^1^

Although *N. gonorrhoeae* can be treated with several classes of antibiotics, including β-lactams, the macrolide azithromycin, and the third-generation cephalosporin ceftriaxone, its remarkable genetic plasticity and natural competence for transformation have contributed to the rapid emergence and global spread of antimicrobial resistance (AMR).^2,3^ Resistant *N. gonorrhoeae* strains were detected within a decade of introducing new treatment regimens. Currently, the last line of defense against gonorrhea involves dual therapy with ceftriaxone and azithromycin. However, resistance to azithromycin has risen significantly over the past five years, a trend that was further accelerated during the COVID-19 pandemic as a result of widespread and often inappropriate antibiotic use.^1,4^

In light of the growing threat posed by AMR, the discovery of new chemical scaffolds with antibacterial activity has become a critical priority. One promising approach involves targeting essential bacterial metabolic pathways that are not exploited by current antibiotics.^5,6^ Among these, undecaprenyl diphosphate synthase (UPPS), a key enzyme in the bacterial isoprenoid biosynthesis pathway, has emerged as a particularly attractive target due to its essential role in cell wall biogenesis. UPPS catalyzes the sequential condensation of eight molecules of isopentenyl pyrophosphate (IPP) with farnesyl pyrophosphate (FPP) to produce undecaprenyl pyrophosphate (UPP), a C55 lipid carrier required for peptidoglycan precursor translocation and the biosynthesis of other essential cell envelope components. Inhibiting this enzyme compromises cell wall integrity and is lethal to many bacteria.^7,8^

UPPS is highly conserved across diverse bacterial pathogens but absent in humans, making it an appealing target for selective antibacterial intervention. Its functional importance has been demonstrated in *Bacillus subtilis*, where reduced UPPS activity sensitized cells to several cell wall–active antibiotics, including fosfomycin and D-cycloserine.^8^ The first class of UPPS inhibitors to be described were bisphosphonates, as reported by Zhu and Guo.^9,10^ These anionic compounds disrupted cell wall synthesis and demonstrated antibacterial activity against Gram-positive organisms. However, they lacked efficacy against Gram-negative bacteria, likely due to poor outer membrane permeability and reduced cellular uptake.^9^ More recently, other scaffolds such as phthalic acid–based inhibitors (e.g., **BPH-1100**) have been identified and structurally characterized in complex with UPPS.^9,11^ While promising, many of these compounds suffer from suboptimal pharmacokinetics. Therefore, continued efforts to discover and optimize drug-like UPPS inhibitors remain essential, particularly for high-priority Gram-negative pathogens such as *N. gonorrhoeae*.

To accelerate the discovery of novel chemical matter targeting *N. gonorrhoeae* UPPS (*Ng*UPPS), we employed weak affinity chromatography coupled to mass spectrometry (WAC-MS)^12^ to screen a structurally diverse fragment library for potential binders. Top-ranking fragments identified by WAC-MS were subsequently soaked into apo-*Ng*UPPS crystals, yielding 16 co-crystal structures that revealed fragment binding along distinct regions of the enzyme’s active site. In addition, we report the co-crystal structure of *Ng*UPPS bound to **BPH-1100**, a known inhibitor scaffold. Together, these structures offer critical insights into ligand-binding hotspots within the active site and provide a valuable framework for future structure-based drug design efforts against this essential and underexploited antimicrobial target.

## Results and Discussion

### Identification of fragments that bind to *Ng*UPPS using WAC-MS

The peptidoglycan biosynthesis pathway has been a major focus for developing antimicrobial agents targeting its later stages. This pathway has led to the development of key inhibitors like vancomycin, methicillin, and bacitracin, which disrupt bacterial cell wall synthesis.^13^ Since mammalian cells lack peptidoglycan, several enzymes involved in this pathway, including UPPS, are considered attractive targets for novel antimicrobial development. To identify *Ng*UPPS ligands from a fragment library, we utilized weak affinity chromatography coupled with mass spectrometry (WAC-MS), a high-throughput, label-free technique known for its sensitivity and ability to detect weak and transient binding interactions.^12^

To enable fragment discovery via WAC-MS, we expressed and purified recombinant *Ng*UPPS from *Escherichia coli* (**Supplementary Figure S1**). The enzymatic activity of the recombinant protein was confirmed using a couple assay with pyrophosphatase (**Supplementary Figure S2)**. Briefly, UPPS catalyzes the addition of up to 8 isopentenyl pyrophosphate (IPP) units to a farnesyl pyrophosphate (FPP) molecule, producing 8 moles of inorganic pyrophosphate. Then, pyrophosphatase hydrolyzes the pyrophosphate inorganic phosphate, which is quantified using Biomol Green reagent.^14^

For WAC-MS experiments, approximately 1 mg of purified *Ng*UPPS was immobilized onto derivatized diol-silica via reductive amination. This process involves covalent coupling between lysine residues on the protein and aldehyde groups on the silica matrix. The immobilization was carried out in batch mode, ensuring proper mixing to maximize protein binding. Following immobilization, enzymatic activity was assessed and found to be comparable to that of the protein in solution **(Supplementary Figure S2C)**. The silica containing immobilized *Ng*UPPS was packed into PEEK tubing and connected to a high-performance liquid chromatography (HPLC) system coupled to a mass spectrometer. A reference column functionalized with ethanolamine (instead of the protein) on diol-silica was prepared in parallel.

In WAC-MS, fragments are separated based on their relative affinities for the immobilized protein target, enabling identification of binders by comparing their retention times to those on a reference column lacking the protein. Using *Ng*UPPS- conjugated silica and the matched empty reference column, we screened a custom fragment library from Key Organics (BIONET®) plus nine carboxylic acid–derived fragments from our internal compound collection, comprising 1,543 fragments that follow the Rule of Three.^15^ The library was organized into pools designed to minimize overlapping mass spectra and maximize performance based on the resolution of the mass spectrometer used. In positive ion mode, compounds within each pool were selected to differ by at least 2 Da in their molecular masses, while in negative ion mode, a minimum difference of 1 Da was maintained. Each pool was injected sequentially, and fragments were identified by their molecular ions, detected as [M+H]⁺ or [M–H]⁻ in positive and negative ionization modes, respectively, using mass spectrometry.

To optimize detection, pooled library runs were first performed on the reerence (no-protein) column using both positive and negative ionization modes, enabling fine-tuning of MS parameters for maximal fragment coverage. Using this strategy, 99.5% of the library was successfully detected, with only 7 out of 1,543 fragments not observed. Retention times on the reference column were below 0.5 minutes, indicating minimal interaction with the silica matrix.

In contrast, when the library was applied to the column containing immobilized *Ng*UPPS, a subset of fragments showed substantial retention shifts. Specifically, 490 fragments exhibited specific retention times (tspec)—defined as the difference in retention time between the *Ng*UPPS and reference columns—ranging from 0.5 to 18.5 minutes. The remaining 1,046 fragments had tspec values below 0.5 minutes, suggesting limited or no interaction with the protein (**Figure 1**). To identify potential hits, a retention time shift of ≥5 minutes was used as an empirical threshold for positive binding. Based on this criterion, 23 putative *Ng*UPPS binders were identified, representing a chemically diverse set of fragments (**Figure 1**, **Supplementary Table S1**).

**Figure 1.**
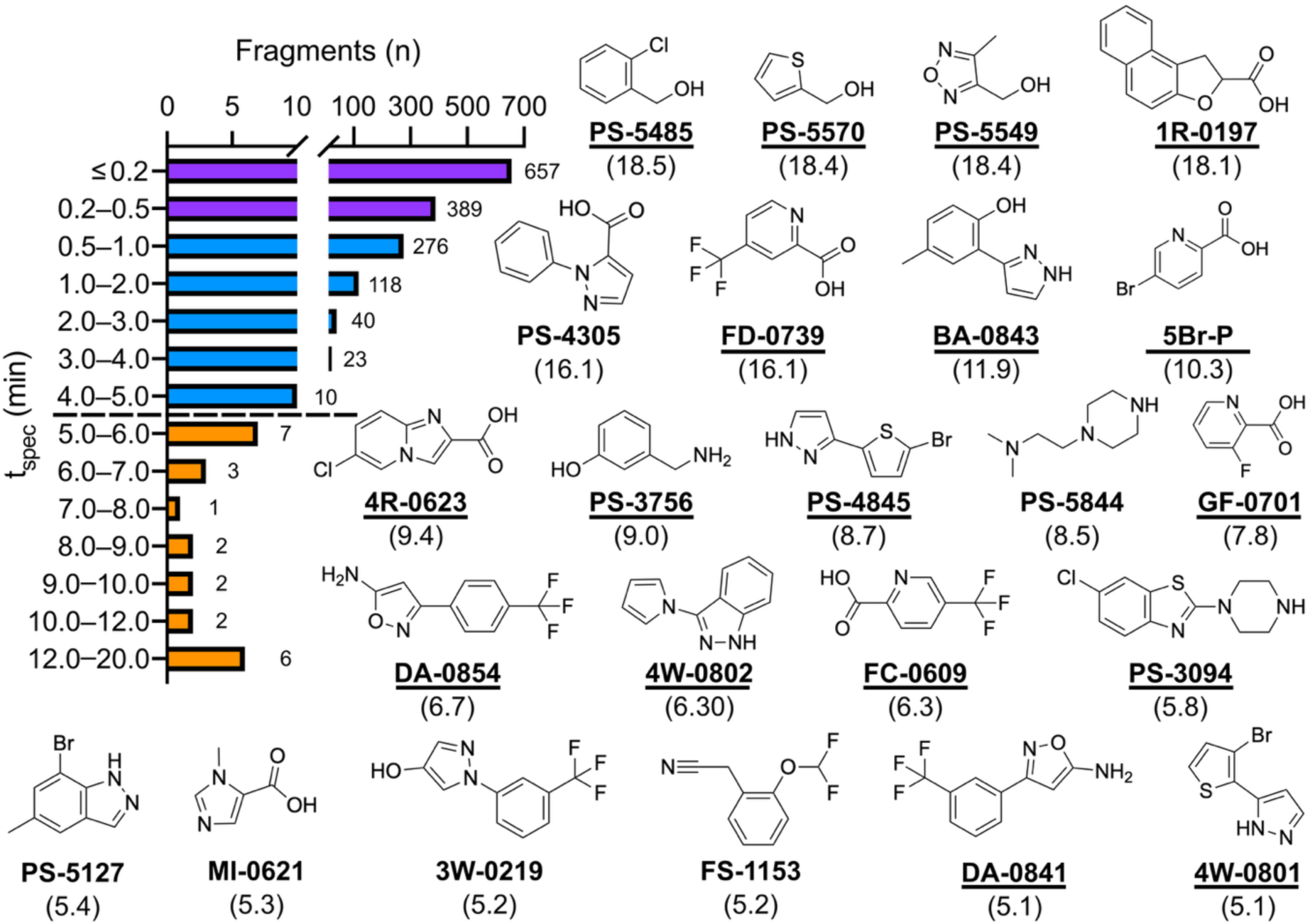
Silica-immobilized *Ng*UPPS retains enzymatic activity and enables fragment interaction profiling. Bar graph showing the distribution of specific retention time shifts (tspec) for 1,537 fragments screened using the *Ng*UPPS affinity column. Numbers above each bar indicate the number of fragments in that bin. Fragments with minimal retention (tspec < 0.5 min) are shown in purple, and weakly retained fragments (0.5 < tspec < 5 min) in blue. A threshold of tspec ≥ 5 min (dashed line) was used to define putative binders (orange). Structures of fragments above this threshold are shown, with retention times (in parentheses) representing averages from two independent experiments. Fragments for which co-crystal structures were determined are underlined. Complete data are provided in Supplementary Table S1.

As expected for fragment-based screening, the identified ligands are not anticipated to exhibit high-affinity binding.^12^ Nevertheless, we attempted to validate these putative hits using our enzymatic activity assay. These efforts were unsuccessful, likely due to a combination of weak fragment binding affinities and the use of high substrate concentrations (15 µM FPP and 120 µM IPP) in the assay, which would outcompete low-affinity fragments for the active site. This highlights a common limitation of fragment screening against active-site enzymes and underscores the utility of WAC-MS for detecting low-affinity binders that may be missed in conventional biochemical assays.

### Crystallographic Analysis of *Ng*UPPS: Apo Structure and Complex with **BPH-1100**

While WAC-MS is highly sensitive to weak binders, it does not inherently provide detailed structural insights into how fragments interact with target proteins. To address this limitation, we integrated WAC-MS with protein crystallography, which not only confirmed fragment binding but also revealed precise binding modes and key protein–ligand interactions at atomic resolution, guiding subsequent optimization through structure-based design.

Although several high-resolution structures of UPPS from various bacterial species have been deposited in the Protein Data Bank (PDB), no structural information was available for *Ng*UPPS at the outset of this study. To overcome this gap, we developed a robust crystallization system for *Ng*UPPS suitable for fragment soaking experiments.

Developing a crystal system amenable to fragment soaking required optimization of several parameters, including crystallization conditions, seeding protocols, and solvent tolerance.^16,17^ Through iterative refinement, we established a system that reliably produced high-quality *Ng*UPPS crystals with diffraction resolutions of approximately 2.0 Å and tolerance to up to 10% DMSO, making them suitable for fragment-based studies.

Using these crystallization conditions, we solved the structure of apo- *Ng*UPPS to 1.6 Å resolution via molecular replacement, employing the *E. coli* UPPS structure (PDB ID: 1JP3; 53% amino acid sequence identity)^18^ as the search model (**Figure 2**; **Supplementary Table S2**). Both *E. coli* and *N. gonorrhoeae* UPPS are Z-type isoprenyl-diphosphate synthases, sharing the prototypical fold first observed in *Micrococcus luteus*.^19^ *Ng*UPPS adopts a characteristic α/β fold distinct from the canonical E-type isoprenoid synthases. The catalytic domain consists of six parallel β-strands surrounded by α-helices, forming a cone-shaped central β-sheet that encloses a large hydrophobic tunnel responsible for product elongation.

**Figure 2.**
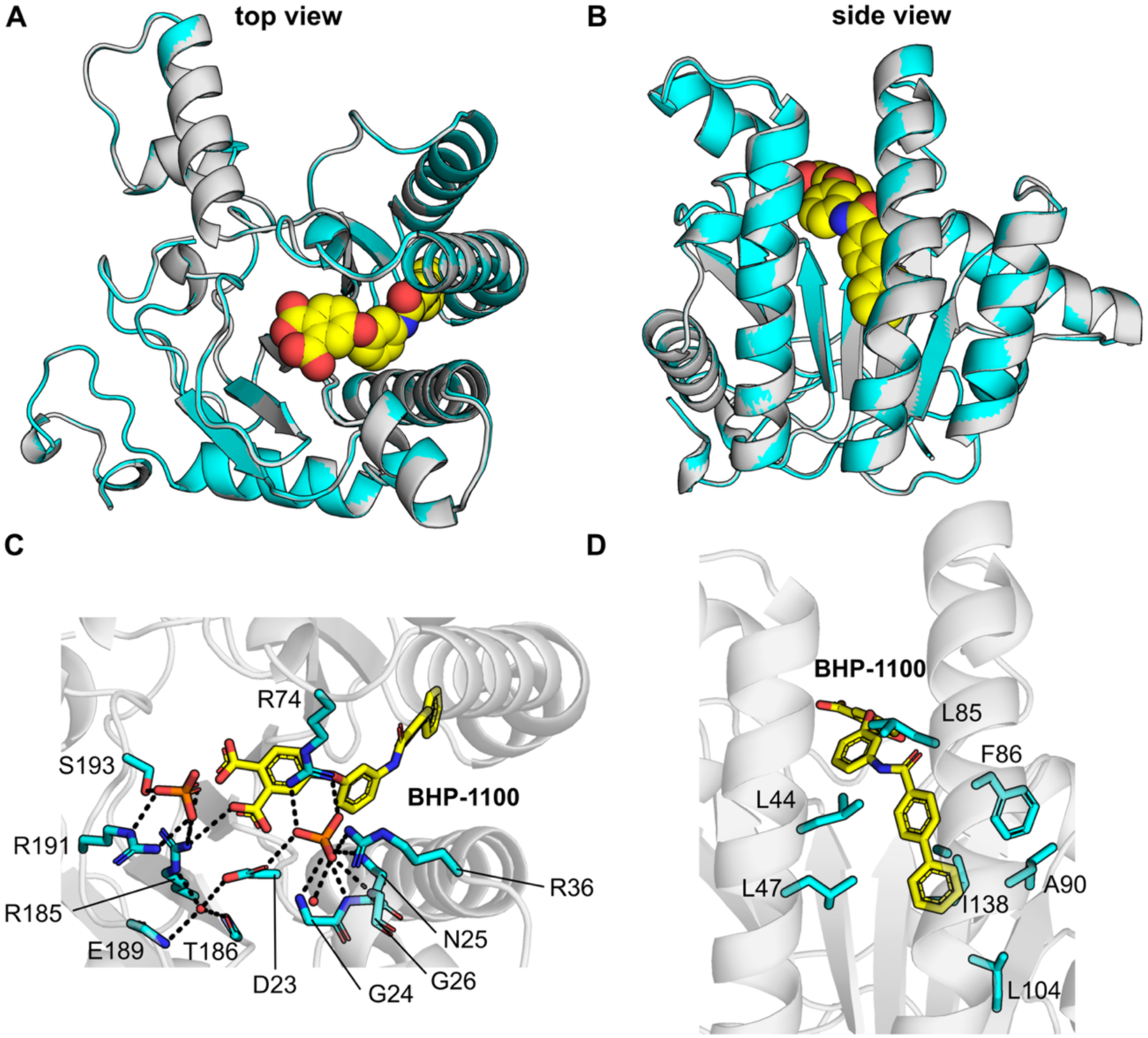
Crystal structures of *Ng*UPPS, apo- and bound to **BHP-1100**. (A, B) Cartoon representations of the apo-*Ng*UPPS structure (cyan; PDB ID: 9OH8) and the **BPH-1100**–bound structure (gray; PDB ID: 9OHA), with the ligand shown as spheres. Structures superimposed using PyMol. (C, D) Binding details of **BPH-1100** within the *Ng*UPPS substrate-binding tunnel. Protein residues (carbon atoms in cyan), ligand (carbon atoms in yellow) and phosphate ions are shown as sticks. Water molecules are shown as red spheres. Dashed black lines represent potential hydrogen bonds.

Apo structures of UPPS have been reported for *E. coli* (PDB IDs: 1JP3, 3QAS)^20,21^, *Mycobacterium tuberculosis* (PDB ID: 2VG4)^22^, and *Helicobacter pylori* (PDB ID: 2D2R)^23^, sharing 53%, 42%, and 40% sequence identity with *Ng*UPPS, respectively. Despite their high structural similarity (overall Cα root mean square deviations - r.m.s.d.; < 1.0 Å), analysis revealed a notable difference in the segment spanning residues 65 to 86 and 110 to 124 (numbering for *Ng*UPPS) (**Supplemental Figures S3A, S3B**). In *E. coli* and *H. pylori* apo-UPPS structures, this segment adopts a conformation distinct from that observed in *N. gonorrhoeae* and *M. tuberculosis* apo-UPPS. This region, located near the tunnel entrance, contributes to substrate binding and catalysis, and likely acts as a hinge to facilitate structural transitions during the elongation cycle.^21^ The conformations observed in *E. coli* and *H. pylori* are consistent with a "closed" state, whereas those of *M. tuberculosis* and *N. gonorrhoeae* represent an "open" state, suggesting a conformational gating mechanism for regulating product elongation and release.^21^

We also determined the co-crystal structure of *Ng*UPPS in complex with **BPH- 1100** (compound **10** in Ref. 10)^10^ at a resolution of 1.8 Å (**Figure 2; Supplementary Table S2**). **BPH-1100** is a phthalic acid–based inhibitor previously reported to have an IC50 of 3.2 µM against *E. coli* UPPS.^10^ Using our enzymatic assay, we determined the IC50 of **BPH-1100** against *Ng*UPPS to be 3.4 µM (**Supplementary Figure S4**). Structural comparison between the apo and **BPH-1100**–bound *Ng*UPPS forms revealed no significant conformational changes (overall Cα r.m.s.d. < 0.2 Å; **Figures 2A and 2B**), indicating that the apo structure is competent for ligand binding.

In the co-crystal structure, the phthalic acid moiety of **BPH-1100** engages in an extensive hydrogen-bonding network with polar side chains of conserved residues (Asp23, Asn25, Arg36, Arg74, Arg185, Thr186, Arg191, and Ser193), the main chain atoms of Gly24, Asn25, and Gly26, coordinated water molecules, and phosphate ions from the solvent (**Figure 2C**). At the opposite end of the ligand, the biphenyl moiety is anchored within a hydrophobic pocket formed by conserved residues (Leu44, Leu47, Leu85, Phe86, Ala89, Leu90, Leu104, and Ile138) whose side chains project into the interior of the substrate-binding tunnel (**Figure 2D**).

The *E. coli* UPPS enzyme has also been previously co-crystallized with **BPH- 1100**.^10^ While the biphenyl moiety occupies a similar position within the tunnel in both the *E. coli* and *N. gonorrhoeae* UPPS structures, the orientation of the phthalic acid group differs markedly at the tunnel entrance (**Supplementary Figures S3C, S3D**). This difference is likely attributable to the *E. coli* structure being captured in a closed conformation, in contrast to the open conformation observed in the *Ng*UPPS– **BPH-1100** complex. Additionally, the *E. coli* structure accommodates a second **BPH-1100** molecule between the two α-helices forming part of the tunnel wall, whereas only a single **BPH-1100** molecule is observed in the *Ng*UPPS structure.

### Structural Mapping of Fragment Binding Across the UPPS Active Site

Having established a robust crystallization system for *Ng*UPPS and validated its ligand-binding competence through co-crystallization with **BPH-1100**, we next sought to characterize fragment binding across the active site. To this end, apo *Ng*UPPS crystals were soaked with 23 fragment hits identified by WAC-MS, and structural analysis was performed to map their binding modes.

Apo-*Ng*UPPS crystals were soaked with 10 mM of each fragment for up to 12 hours at 24 °C, using a maximum DMSO concentration of 10%. Crystals were harvested and cryoprotected in mother liquor supplemented with 15% glycerol. X-ray diffraction data were collected and processed individually. Of the 23 fragments tested, 15 yielded high-quality co-crystal structures with resolutions ranging from 1.4 to 2.3 Å (**Supplementary Figure S5A, S5B**; **Supplementary Table S2**). These structures were isomorphous with both the apo- and **BPH-1100**–bound *Ng*UPPS forms, crystallizing in the C2 space group with closely matching unit cell dimensions (deviations in cell lengths < 0.4 Å and angles < 0.3°). For one additional fragment, soaking was unsuccessful; however, co-crystallization yielded well-diffracting crystals in the P21 space group (resolution of 2.3 Å) that retained a fold highly similar to the apo- and **BPH-1100**–bound structures (**Supplementary Figure S5C, S5D**). Across all fragment-bound complexes, the final refined models were virtually identical to the apo structure, with overall Cα r.m.s.d. values of < 0.3 Å, confirming structural integrity and suitability for confident interpretation of fragment binding.

Because weakly bound fragments often exhibit partial occupancy, ligand assessment was conducted following full refinement of the protein backbone, side-chain rotamers, and water molecules. Fragments were modeled at fixed occupancies ranging from 0.25 to 1.0 and were validated using Polder omit maps (Fo–Fc) and local real-space correlation coefficients (CC)^24^.

Among the 15 fragment-bound structures that crystallized in the C2 space group (each containing a single copy of *Ng*UPPS in the asymmetric unit), 11 displayed a single binding site with a single well-defined ligand conformation. In three structures, the fragment occupied two distinct sites, each with a single conformation (**PS-5549**, **4R-0623** and **GF-701**). In the remaining C2 structure, the fragment was observed at two sites: one with a single conformation and the other showing three alternate conformers (**PS-5570**), indicative of local binding flexibility. In the structure crystallized in the P21 space group (**1R-0197**), two copies of *Ng*UPPS were present in the asymmetric unit; in one protomer, the fragment bound in a single pose, while in the second protomer, two conformers of the fragment occupied the same binding site (**Supplementary Figures S6, S7, S8**).

These high-resolution fragment-bound structures provided a detailed spatial framework for mapping ligand binding across the *Ng*UPPS active site. To facilitate interpretation, the substrate-binding tunnel was conceptually divided into four regions following the framework proposed by Zhu and colleagues^10^: regions 1 and 2 (top) correspond to the pyrophosphate-binding pockets near the tunnel entrance; region 3 (middle) encompasses the central hydrophobic core; and region 4 (bottom) spans the distal end of the tunnel, where the elongating isoprenoid chain is accommodated (**Figure 3**).

**Figure 3.**
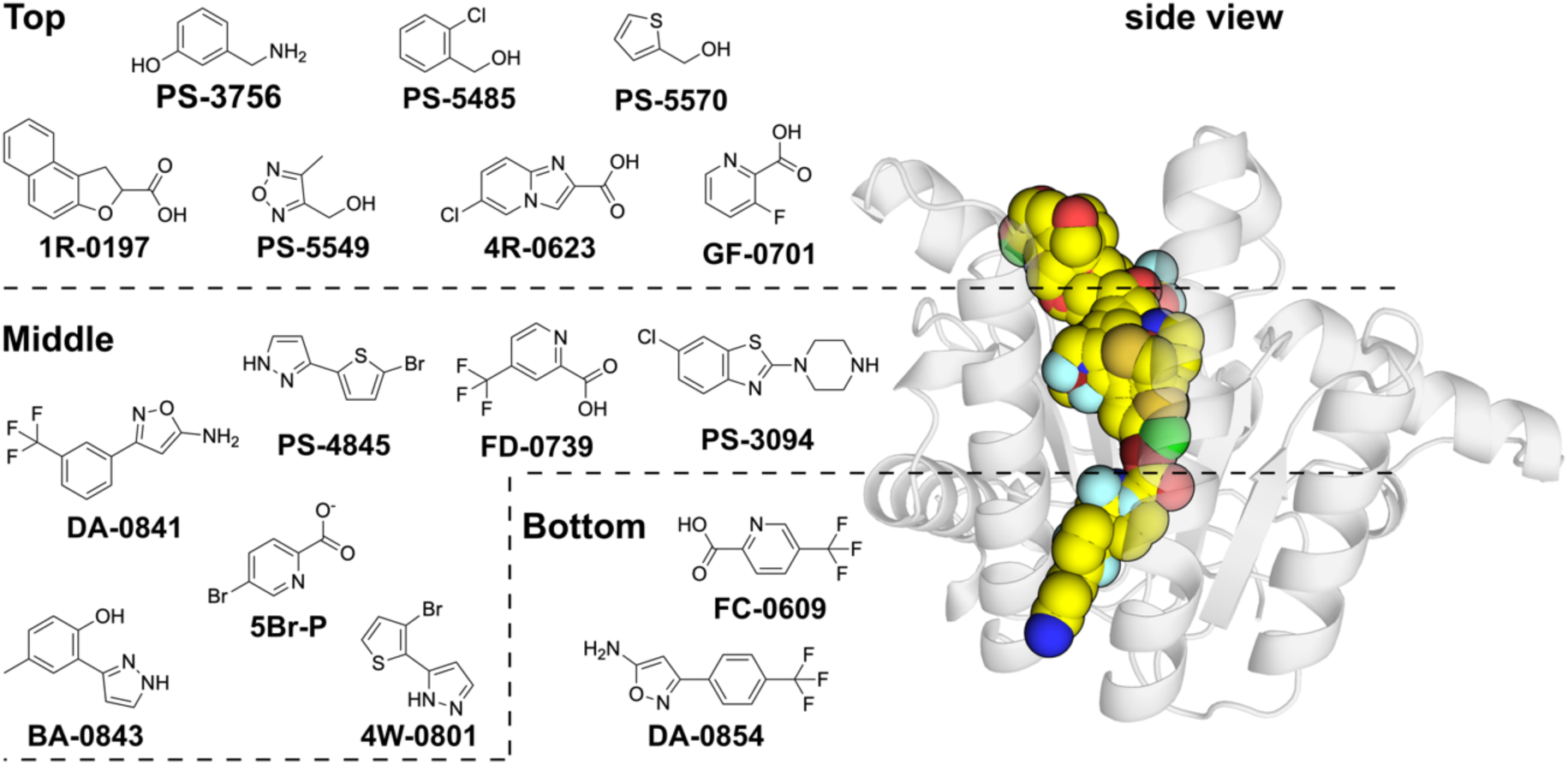
Mapping of the 16 identified fragments onto the *Ng*UPPS substrate-binding site. The protein is shown as a semi-transparent gray cartoon. Fragments occupying the binding site are depicted as spheres. Mapping was performed by aligning all fragment-bound structures to the *Ng*UPPS–**4R-0623** complex, which is shown as a cartoon for reference. PDB IDs of apo-*Ng*UPPS and NgUPPS bound to fragments are: 9OH8 (Apo) 9OI4 (**1R-0197**), 9OHI (**PS-5549**), 9OHJ (**4R-0623**), 9OIW (**GF-0701**), 9OHO (**PS-3756**), 9OHB (**PS-5485**) and 9OHG (**PS-5570**), 9OJE (**5Br-P**), 9OI6 (**FD-0739**), 9OIY (**DA-0841**), 9OIV (**4W-0801**), 9OJB (**PS-0394**), 9OI5 (**PS-4845**), 9OHP (**BA-0843**), 9OJ1 (**DA-0854**), 9OJA (**FC-0609**).

Regions 1 and 2 (top) are located near the entrance of the catalytic tunnel and are characterized by a predominantly positive electrostatic surface, consistent with their role in coordinating the negatively charged pyrophosphate moieties of the natural substrates. In the *E. coli* UPPS structure, region 1 binds the IPP substrate through coordination with a Mg^2+^ ion.^25^ In our *Ng*UPPS fragment-bound structures, seven fragments were observed binding within region 1 (**1R-0197**, **PS-5549**, **4R- 0623**, **GF-0701**, **PS-3756**, **PS-5485** and **PS-5570**) (**Supplementary Figure S6**). Most of these fragments form hydrogen bonds with main chain atoms of Met22, Gly24, Asn25, Gly26, and Arg27. Some fragments, including **PS-5485**, **PS-5570** and **PS-5549**, also engage side chain atoms, particularly those of Arg27, Asn25, and Arg74.

Region 2 lies slightly deeper within the tunnel and maintains a similarly positive electrostatic environment. Fragments identified in this region include fragments **DA-0841**, **FD-0739**, and **5Br-P**. These mostly interact with solvent molecules found in this region of the tunnel and do not engage protein residues directly (**Supplementary Figure S7**).

In contrast, regions 3 (middle) and 4 (bottom) occupy the more distal, solvent-shielded portions of the tunnel and are lined predominantly by hydrophobic residues, reflecting their role in stabilizing the isoprenoid tail during elongation (**Supplementary Figure S8**). Four fragments were identified in region 3 that primarily exploit van der Waals and hydrophobic interactions: **PS-3094**, **PS-4845**, **BA-0843**, and **4W-0801**. Notably, fragment **4W-0801** also forms a hydrogen bond with the side chain of Asn25.

Region 4, located at the distal end of the tunnel, has previously been shown to accommodate a second copy of substrate-mimetic inhibitors such as **BPH-629** (PDB ID: 2E98) and **BPH-1100** (PDB ID: 3SGX) in *E. coli* UPPS.^10^ Fragments **FC-0609** and **DA-0854** were observed binding in this region.

Together, the crystallographic analysis demonstrates that the fragment hits span all four functional regions of the *Ng*UPPS substrate-binding tunnel, from the pyrophosphate-coordinating entrance to the distal hydrophobic pocket. This comprehensive spatial coverage highlights the tractability of the UPPS active site and provides a strong framework for fragment-based drug design. The diversity of binding modes—ranging from ionic interactions with basic residues to deep hydrophobic contacts—offers multiple opportunities for fragment merging, linking, or growing strategies. These structures represent a critical first step toward the development of potent UPPS inhibitors targeting drug-resistant *N. gonorrhoeae*.

## Conclusions

The rapid emergence of antimicrobial resistance in *N. gonorrhoeae* underscores the urgent need for new therapeutic strategies targeting essential bacterial pathways not exploited by current antibiotics. Here, we demonstrate a comprehensive pipeline for the identification and structural characterization of fragment binders targeting *Ng*UPPS, a critical enzyme in bacterial cell wall biosynthesis.

Using weak affinity chromatography coupled to mass spectrometry (WAC- MS), we screened a diverse fragment library and identified 23 putative *Ng*UPPS binders. Follow-up structural studies successfully yielded 16 fragment-bound crystal structures, revealing ligand engagement across four distinct regions of the enzyme’s hydrophobic tunnel. This broad spatial coverage, encompassing both the pyrophosphate-binding site and the distal hydrophobic pocket, highlights the high ligandability of the *Ng*UPPS active site and provides a structural basis for rational fragment merging, linking, and growing strategies.

Additionally, we determined the high-resolution structure of *Ng*UPPS bound to **BPH-1100**, a phthalic acid–based inhibitor previously characterized against *E. coli* UPPS. Comparison of apo, fragment-bound, and inhibitor-bound structures revealed that the *Ng*UPPS active site adopts an open conformation competent for ligand binding without requiring substantial conformational rearrangements. Notably, conserved polar and hydrophobic residues lining the tunnel contribute to the binding of both polar and nonpolar fragments, suggesting that mixed pharmacophore designs may be advantageous for inhibitor development.

Together, these findings establish the structural tractability of *Ng*UPPS for drug discovery and provide critical starting points for the development of next-generation inhibitors targeting this essential enzyme. Our integrated WAC-MS and crystallographic approach serves as a powerful platform for fragment-based discovery efforts against challenging antimicrobial targets.

## Methods

### Cloning and recombinant production of *Ng*UPPS

Genomic DNA from *Neisseria gonorrhoeae* strain NCTC 13822 was purified using the DNeasy Plant Mini Kit (QIAGEN), following the manufacturer’s instructions.

*N. gonorrhoeae* NCTC 13822 is a multidrug-resistant clinical isolate, exhibiting resistance to azithromycin, low-level resistance to ceftriaxone, high-level resistance to cefixime, and resistance to penicillin G and tetracycline.^26^ The integrity of the purified genomic DNA was assessed by agarose gel electrophoresis and the DNA was stored at -20 °C for further analysis.

The full-length *uppS* gene from *Neisseria gonorrhoeae* strain NCTC 13822 (encoding residues Met1 to Asn248) was amplified by PCR and cloned into the pNIC-Bsa4 vector using ligation-independent cloning (LIC). The primers used were: forward 5′-TACTTCCAATCCATGGGTTTTCTGCAAGGCAAAAAAATTCT-3′ and reverse 5′-TATCCACCTTTACTGTCATCCCTCGGTGCTCAAGGC-3′. Cloning into pNIC-Bsa4 resulted in the addition of an N-terminal 6×His tag followed by a TEV protease cleavage site. Positive clones were verified by sequencing to confirm the absence of mutations and subsequently transformed into *E. coli* BL21(DE3) pRARE cells for protein expression.

A single colony was first inoculated into 50 mL Luria–Bertani (LB) broth supplemented with 50 µg/mL kanamycin and 34 µg/mL chloramphenicol, and incubated at 37 °C with shaking at 130 rpm for 14 h. The overnight culture was then used to inoculate 1.5 L of Terrific Broth (TB) containing 50 µg/mL kanamycin. Protein expression was induced with 200 µM IPTG when the culture reached an OD600 of 1.2. Upon induction, the temperature was reduced to 16 °C and incubation continued at 130 rpm for 16 h. Cells were harvested by centrifugation, suspended in 2x Lysis (1 mL of buffer to 1 gram of cell pellet - wet weight) and stored at -80 °C. 2x Lysis buffer is: 80 mM HEPES (pH 7.5), 600 mM NaCl, 20% glycerol, 1.0 mM TCEP, 20 mM imidazole, and 2 mM PMSF. Cells were lysed by sonication using a Vibracell VCX750 sonicator (Sonics®) set to 35% amplitude with 5-second on/10-second off cycles. Cell debris was removed by centrifugation at 40,000 × g for 45 minutes at 4 °C.

The resulting clarified lysate was loaded onto a 5 mL HisTrap FF crude column (Cytiva®) using an ÄKTA Pure system at a flow rate of 2 mL/min. The column was washed with 10 column volumes of wash buffer (same as lysis buffer but containing 50 mM imidazole), and bound protein was eluted with elution buffer containing 300 mM imidazole. Eluted fractions were pooled and treated with His-tagged TEV protease at a 1:20 molar ratio (*Ng*UPPS:TEV), followed by overnight dialysis at 4 °C against gel filtration buffer (20 mM HEPES, pH 7.5, 300 mM NaCl, 5% glycerol, 0.5 mM TCEP).

The following day, the dialyzed solution was passed through 4 mL of Ni²⁺- charged chelating Sepharose resin to remove TEV protease and any uncleaved His-tagged *Ng*UPPS. The flow-through, containing cleaved *Ng*UPPS, was collected, concentrated, and subjected to size-exclusion chromatography using a Superdex 200 pg 16/600 column for final polishing. Fractions corresponding to *Ng*UPPS were pooled and concentrated to 12 mg/mL.

### *Ng*UPPS immobilization in silica beads and column preparation

Purified *Ng*UPPS was covalently immobilized onto diol-functionalized spherical silica gel (300 Å pore size, 5 µm particle size, endcapped; Sorbtech, Norcross, GA). Reagents including HPLC-grade ammonium acetate, dimethyl sulfoxide (DMSO), ethanolamine, 4-(2-hydroxyethyl)piperazine-1-ethanesulfonic acid (HEPES), periodic acid, and sodium cyanoborohydride were purchased from Sigma-Aldrich.

Immobilization was performed in batch mode following the protocol described by Ohlso and Duong-Thi.^12^ Briefly, for each batch, 9 mg of diol silica was suspended in 250 µL of isopropanol in a 1.5 mL microcentrifuge tube, sonicated for 2 minutes, and centrifuged at 4,000 × g for 2 minutes. This was followed by three washes with 250 µL of ultrapure water. The silica was then suspended in 250 µL of an aqueous 100 mg/mL periodic acid solution and incubated at 22 °C for 2.5 hours with shaking at 1,000 rpm on a ThermoMixer™ C (Eppendorf). After oxidation, the silica was washed three times with ultrapure water and three times with gel filtration buffer (250 µL per wash).

To prepare the columns, the activated silica was incubated with either protein solution (for target columns) or buffer alone (for reference columns). For reference columns, oxidized silica was incubated with 250 µL of 1 M ethanolamine containing 10 mg/mL sodium cyanoborohydride to block unreacted aldehyde groups. The target column was prepared using 1 mL of *Ng*UPPS at a concentration of 1 mg/mL, with all sodium cyanoborohydride-containing solutions maintained at 10 mg/mL. Following protein coupling, the immobilization reaction was terminated by washing the silica four times with 250 µL of gel filtration buffer. Supernatants and wash fractions were collected, and absorbance at 280 nm was measured to indirectly estimate the immobilization yield. Residual aldehyde groups were quenched by incubation with 1 M ethanolamine containing 10 mg/mL sodium cyanoborohydride for 2.5 hours at 22 °C with shaking at 500 rpm. The *Ng*UPPS-bound silica was then washed three additional times with gel filtration buffer (250 µL each) and stored in the same buffer supplemented with 0.1% sodium azide until use.

For column packing, both *Ng*UPPS-bound and reference silica were slurry-packed into 4 cm PEEK tubing (1/16", 500 μm i.d.) using gel filtration buffer and an HPLC pump (1260 Infinity II Quaternary Pump, Agilent) operating at 300 bar for 1 hour. Each tubing was fitted with a zero-dead-volume union containing a glass fiber frit to retain the packing material and serve as the analytical column housing.

A ThermoMixer™ C (Eppendorf) was used for mixing and incubation during the protein immobilization steps. Protein concentration in supernatants and washes was monitored using a NanoDrop 2000 spectrophotometer (Thermo Scientific) to assess immobilization efficiency. A zero-dead-volume union (1/16″, 0.25 mm bore) was obtained from VICI Valco®. Glass microfiber filters (GF/A, Whatman™) and PEEK tubing (1/16″ outer diameter, 500 µm inner diameter) were purchased from GE Healthcare.

### Screening of fragments library

The fragment library used in this study comprised 1,534 compounds sourced from the Key Organics BIONET Fragment Library. Fragments were organized into mixtures containing 12 compounds each, prepared at 50 µM per fragment in 20 mM ammonium acetate with 2% DMSO. To ensure mass resolution during MS detection, mixtures were assembled such that each fragment differed by at least 2 Da in positive ion mode and 1 Da in negative ion mode. Additionally, nine hand-picked carboxylic acid–derived fragments were included, bringing the total library size to 1,543 compounds.

Fragment screening by weak affinity chromatography (WAC) was performed using an Agilent 1260 Infinity II LC system equipped with either the *Ng*UPPS-bound (target) column or the reference column (no protein). Fragment mixtures (0.5 µL) were injected in duplicate under isocratic elution with 20 mM ammonium acetate at a flow rate of 15 µL/min and 25 °C. Detection was performed using a single quadrupole mass spectrometer (Agilent InfinityLab LC/MSD) operated with an electrospray ionization (ESI) source.

The initial screen was conducted in positive ion mode (ESI+), with signal acquisition in selected ion monitoring (SIM) mode targeting [M+H]+. The total run time per injection was 26 minutes. Fragments not detected under ESI+ conditions were reanalyzed in negative ion mode (ESI−), targeting [M−H]−. ESI source parameters were as follows: capillary voltage 3,200 V (ESI+) and 2,200 V (ESI−), nebulizer pressure 30.0 bar (N2), dry gas flow 8.0 L/min at 300 °C, fragmentor voltage 110 V (ESI+) and 100 V (ESI−), and quadrupole temperature maintained at 100 °C.

Specific retention time (tspec) was calculated using the formula:

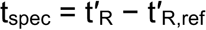

where t′R and t′R,ref are the adjusted retention times for the target and reference columns, respectively, obtained by subtracting the retention time of the void marker (DMSO) from the observed retention time (tR):

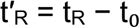

Between analyses, columns were stored at 4 °C in gel filtration buffer containing 0.1% sodium azide.

### *Ng*UPPS crystallization and structure determination

*Ng*UPPS was concentrated to 12 mg/mL (422 µM) and screened for crystallization using sitting-drop vapor diffusion with commercially available solution sets from Molecular Dimensions. Drops were set up using a Mosquito liquid handler (SPT Labtech). The best-diffracting crystals were obtained at 20 °C in a condition containing 0.056 M sodium phosphate monobasic monohydrate and 1.344 M potassium phosphate dibasic, pH 8.2. Crystals with the highest resolution (1.7– 1.9 Å) were consistently observed at a 1:1 volume ratio of protein to crystallization solution, and this condition was subsequently used for fragment soaking experiments.

For co-crystallization with **BPH-1100** (compound **10** in Ref. 10) [4-{3- [(biphenyl-4-ylcarbonyl)amino]phenoxy}benzene-1,2-dicarboxylic acid; Cat# SKT098506, Vitas-M Laboratory], *Ng*UPPS was concentrated to 12 mg/mL and incubated with 3 mM of the compound (a 7-fold molar excess) for 1 hour at room temperature prior to crystallization setup. Co-crystals were obtained using sitting-drop vapor diffusion in a condition containing 30% PEG 4000, 0.1 M Tris-HCl (pH 8.5), and 0.2 M lithium sulfate, employing a 1:2 volume ratio of crystallization solution to protein.

### DMSO tolerance assay and fragments soaking

To determine the maximum DMSO concentration compatible with fragment soaking, pre-formed *Ng*UPPS crystals were exposed to increasing concentrations of DMSO (ranging from 2% to 30%) to identify the highest concentration that caused no visible crystal damage and preserved high-resolution X-ray diffraction. Based on this evaluation, 10% DMSO was selected as the optimal soaking condition.

Selected fragments were diluted to 10 mM in crystallization buffer, and 2 µL of this solution was gently pipetted onto the crystallization drop. Crystals were incubated for 12 hours at room temperature. Prior to cryocooling, the soaking solution was supplemented with glycerol to a final concentration of 10% for cryoprotection. Crystals were mounted in nylon loops (MiTeGen) and flash-frozen in liquid nitrogen for data collection.

X-ray diffraction data were collected at 100 K at the Sirius Manacá beamline (24-ID-E) at the Brazilian Synchrotron Light Laboratory. Data integration and scaling were performed using XDS^27^ and AIMLESS (CCP4 suite),^28^ respectively. Structure determination was carried out by molecular replacement using DIMPLE (CCP4 suite),^28^ employing the *E. coli* UPPS structure (PDB ID: 1JP3; 53% amino acid sequence identity)^18^ as the search template.

Fragment electron density was inspected using WinCoot,^29^ and fragment presence was further confirmed using Polder map calculations,^24^ which were particularly useful for fragments exhibiting low occupancy.

### *Ng*UPPS enzymatic assay

The *Ng*UPPS enzymatic assay was optimized using the following final concentrations: 20 nM *Ng*UPPS, 15 µM farnesyl pyrophosphate (FPP), 120 µM isopentenyl pyrophosphate (IPP), and 0.001 U/µL pyrophosphatase. All components were solubilized in enzymatic buffer containing 50 mM HEPES (pH 7.5), 1 mM MgCl₂, 150 mM NaCl, 0.005% BSA, 0.01% CHAPS, and 1% DMSO to simulate the conditions used in dose–response assays.

Stock solutions of each reagent were pre-diluted to four times the final assay concentration. Then, 2.5 µL of each component was dispensed into non-binding, flat-bottom 384-well plates (Corning #3575). To initiate colorimetric detection of released inorganic phosphate, 20 µL of BioMol® Green reagent (Enzo Biochem) was added to each well. After a 20-minute incubation at room temperature, absorbance was measured at 620 nm using a SpectroStar Nano microplate reader (BMG Labtech).

To determine the half-maximal inhibitory concentration (IC50) of **BPH-1100**, the compound was dissolved in 100% DMSO to prepare a 20 mM stock solution. A 12-point, two-fold serial dilution was prepared in 100% DMSO, starting from 20 mM. To reduce the final DMSO concentration in the assay, a 1:10 pre-dilution of each concentration was made in enzymatic buffer. From this pre-dilution, 1 µL was added to 1.5 µL of *Ng*UPPS, and the mixture was incubated for 20 minutes at room temperature. Separately, a substrate/enzyme mix was prepared containing equal volumes of farnesyl pyrophosphate (FPP), isopentenyl pyrophosphate (IPP), and pyrophosphatase in reaction buffer. Then, 7.5 µL of this mixture was added to the

2.5 µL *Ng*UPPS-compound solution, yielding final assay concentrations of 15 µM FPP, 120 µM IPP, 0.004 U/µL pyrophosphatase, and 20 nM *Ng*UPPS. Following this dilution scheme, the highest inhibitor concentration was 0.2 mM. The reaction was incubated for 40 minutes at 28 °C. To terminate the reaction and develop color, 20 µL of BioMol® Green reagent (Enzo Biochem) was added. After 20 minutes of incubation at room temperature, absorbance was measured at 620 nm. Percent enzyme inhibition was calculated relative to vehicle controls. IC50 values were determined by fitting the dose–response data to a four-parameter logistic (4PL) model using GraphPad Prism (GraphPad Software v. 10.4, San Diego, CA).

## Associated Content

### Supporting Information

Supplementary Figures S1-S8 and Supplementary Tables S1 and S2 (PDF).

### Accession Codes

Atomic coordinates and experimental data have been deposited in the PDB. PDB IDs of apo-NgUPPS and NgUPPS bound to fragments are: 9OH8 (Apo) 9OI4 (**1R-0197**), 9OHI (**PS-5549**), 9OHJ (**4R-0623**), 9OIW (**GF-0701**), 9OHO (**PS-3756**), 9OHB (**PS-5485**) and 9OHG (**PS-5570**), 9OJE (**5Br-P**), 9OI6 (**FD-0739**), 9OIY (**DA-**

**0841**), 9OIV (**4W-0801**), 9OJB (**PS-0394**), 9OI5 (**PS-4845**), 9OHP (**BA-0843**), 9OJ1 (**DA-0854**), 9OJA (**FC-0609**). PDB ID of **BHP-1100**-bound *Ng*UPPS is 9OHA.

## Author Information

### Corresponding authors

**Rafael M. Couñago** - Center of Medicinal Chemistry (CQMED), Center for Molecular Biology and Genetic Engineering (CBMEG), Universidade Estadual de Campinas, UNICAMP, 13083-886-Campinas, SP, Brazil; Structural Genomics Consortium and Division of Chemical Biology and Medicinal Chemistry, UNC Eshelman School of Pharmacy, University of North Carolina, Chapel Hill, North Carolina 27599, USA;

**Katlin B. Massirer** - Center of Medicinal Chemistry (CQMED), Center for Molecular Biology and Genetic Engineering (CBMEG), Universidade Estadual de Campinas, UNICAMP, 13083-886-Campinas, SP, Brazil;

**Gabriela Barreiro** - Eurofarma Laboratórios S/A, 06696-000-Itapevi, SP, Brazil;

### Authors

**André da Silva Santiago** - Center of Medicinal Chemistry (CQMED), Center for Molecular Biology and Genetic Engineering (CBMEG), Universidade Estadual de Campinas, UNICAMP, 13083-886-Campinas, SP, Brazil.

**Bruno Sérgio do Amaral** - Center of Medicinal Chemistry (CQMED), Center for Molecular Biology and Genetic Engineering (CBMEG), Universidade Estadual de Campinas, UNICAMP, 13083-886-Campinas, SP, Brazil.

**Edgar Llomp** - Center of Medicinal Chemistry (CQMED), Center for Molecular Biology and Genetic Engineering (CBMEG), Universidade Estadual de Campinas, UNICAMP, 13083-886-Campinas, SP, Brazil.

**Priscila Zonzini Ramos** - Center of Medicinal Chemistry (CQMED), Center for Molecular Biology and Genetic Engineering (CBMEG), Universidade Estadual de Campinas, UNICAMP, 13083-886-Campinas, SP, Brazil.

**Micael Rodrigues Cunha** - Center of Medicinal Chemistry (CQMED), Center for Molecular Biology and Genetic Engineering (CBMEG), Universidade Estadual de Campinas, UNICAMP, 13083-886-Campinas, SP, Brazil.

**Lucas Rodrigo de Souza** - Center of Medicinal Chemistry (CQMED), Center for Molecular Biology and Genetic Engineering (CBMEG), Universidade Estadual de Campinas, UNICAMP, 13083-886-Campinas, SP, Brazil.

**Erika Piccirilo** - Eurofarma Laboratórios S/A, 06696-000-Itapevi, SP, Brazil.

**Vitor Medeiros Almeida** - Eurofarma Laboratórios S/A, 06696-000-Itapevi, SP, Brazil.

**Ronaldo Aloise Pilli** - Center of Medicinal Chemistry (CQMED); Department of Organic Chemistry, Institute of Chemistry, Universidade Estadual de Campinas, UNICAMP, 13083-970-Campinas, SP, Brazil

### Author Contributions

**A.S.S.** and **B.S.A.** contributed equally to this work.

**B.S.A.** and **L.R.S** designed, executed, and analyzed the WAC-MS experiments.

**A.S.S.** designed, executed, and analyzed the biochemical assays, crystallization experiments, and structure determination.

**E.L.** performed protein structure refinement and managed PDB data deposition.

**P.Z.R.** designed, executed, and analyzed molecular biology and cloning experiments.

**E.P.** conducted the literature review for the project.

**M.R.C.** analyzed the physicochemical properties of the identified fragments.

**V.M.A.** coordinated project activities.

**R.M.C.**, **K.B.M.**, **R.A.P.**, and **G.B.** provided project coordination and secured funding.

**A.dS.S.** and **R.M.C.** wrote the manuscript.

All authors reviewed and approved the final version of the manuscript.

### Notes

The authors declare no competing financial interest.

## Acknowledgments

We thank all members of CQMED-UNICAMP for their help and support. This work was supported by Eurofarma Laboratórios S/A, FAPESP (Fundação de Amparo à Pesquisa do Estado de São Paulo – grant numbers 2013/50724-5, 2014/50897-0, 20/02006-0 and 24/02215-9), Embrapii (Empresa Brasileira de Pesquisa e Inovação Industrial), CNPq (Conselho Nacional de Desenvolvimento Científico e Tecnológico – grant number: 465651/2014-3). P.Z.R. (24/16466-3), M.R.C. (21/04853-4), and L.R.S. (20/16094-8 and 24/09297-0) are recipients of FAPESP fellowships. We thank the staff of the Proteomics section of the Life Sciences Core Facility (LaCTAD), part of the University of Campinas (UNICAMP). This research used facilities of the Brazilian Synchrotron Light Laboratory (LNLS), part of the Brazilian Center for Research in Energy and Materials (CNPEM), a private non-profit organization under the supervision of the Brazilian Ministry for Science, Technology, and Innovations (MCTI). The Manacá beamline staff is acknowledged for the assistance during the experiments (proposal number 231116).

## References

(1) Unemo, M.; Seifert, H. S.; Hook, E. W.; Hawkes, S.; Ndowa, F.; Dillon, J.-A. R. Gonorrhoea. Nat Rev Dis Primers 2019, 5 (1), 79. 10.1038/s41572-019-0128-6.

(2) Rotman, E.; Seifert, H. S. The Genetics of *Neisseria* Species. Annu. Rev. Genet. 2014, 48 (1), 405–431. 10.1146/annurev-genet-120213-092007.

(3) Akinboro, M. K.; Mmaduabuchi, J.; Beeko, P. K. A.; Egwuonwu, O. F.; Oluwalade, O. P.; Akueme, N. T.; Iyioku, B. O.; Okobi, O. E.; Oghenetega, E. P. Epidemiological Trends and Factors Associated With the Morbidity Rate of Gonorrhea: A CDC-WONDER Database Analysis. Cureus 2023. 10.7759/cureus.42981.

(4) Kueakulpattana, N.; Wannigama, D. L.; Luk-in, S.; Hongsing, P.; Hurst, C.; Badavath, V. N.; Jenjaroenpun, P.; Wongsurawat, T.; Teeratakulpisan, N.; Kerr, S. J.; Abe, S.; Phattharapornjaroen, P.; Shein, A. M. S.; Saethang, T.; Chantaravisoot, N.; Amarasiri, M.; Higgins, P. G.; Chatsuwan, T. Multidrug-Resistant Neisseria Gonorrhoeae Infection in Heterosexual Men with Reduced Susceptibility to Ceftriaxone, First Report in Thailand. Sci Rep 2021, 11 (1), 21659. 10.1038/s41598-021-00675-y.

(5) Prasad, N. K.; Seiple, I. B.; Cirz, R. T.; Rosenberg, O. S. Leaks in the Pipeline: A Failure Analysis of Gram-Negative Antibiotic Development from 2010 to 2020. Antimicrob Agents Chemother 2022, 66 (5), e00054–22. 10.1128/aac.00054-22.

(6) Butler, M. S.; Cooper, M. A. Antibiotics in the Clinical Pipeline in 2011. J Antibiot 2011, 64 (6), 413–425. 10.1038/ja.2011.44.

(7) Apfel, C. M.; Takács, B.; Fountoulakis, M.; Stieger, M.; Keck, W. Use of Genomics To Identify Bacterial Undecaprenyl Pyrophosphate Synthetase: Cloning, Expression, and Characterization of the Essential *uppS* Gene. J Bacteriol 1999, 181 (2), 483–492. 10.1128/JB.181.2.483-492.1999.

(8) Lee, Y. H.; Helmann, J. D. Reducing the Level of Undecaprenyl Pyrophosphate Synthase Has Complex Effects on Susceptibility to Cell Wall Antibiotics. Antimicrob Agents Chemother 2013, 57 (9), 4267–4275. 10.1128/AAC.00794-13.

(9) Guo, R.-T.; Cao, R.; Liang, P.-H.; Ko, T.-P.; Chang, T.-H.; Hudock, M. P.; Jeng, W.-Y.; Chen, C. K.-M.; Zhang, Y.; Song, Y.; Kuo, C.-J.; Yin, F.; Oldfield, E.; Wang, A. H.-J. Bisphosphonates Target Multiple Sites in Both *Cis* - and *Trans* -Prenyltransferases. Proc. Natl. Acad. Sci. U.S.A. 2007, 104 (24), 10022– 10027. 10.1073/pnas.0702254104.

(10) Zhu, W.; Zhang, Y.; Sinko, W.; Hensler, M. E.; Olson, J.; Molohon, K. J.; Lindert, S.; Cao, R.; Li, K.; Wang, K.; Wang, Y.; Liu, Y.-L.; Sankovsky, A.; De Oliveira, C. A. F.; Mitchell, D. A.; Nizet, V.; McCammon, J. A.; Oldfield, E. Antibacterial Drug Leads Targeting Isoprenoid Biosynthesis. Proc. Natl. Acad. Sci. U.S.A. 2013, 110 (1), 123–128. 10.1073/pnas.1219899110.

(11) Sinko, W.; Wang, Y.; Zhu, W.; Zhang, Y.; Feixas, F.; Cox, C. L.; Mitchell, D. A.; Oldfield, E.; McCammon, J. A. Undecaprenyl Diphosphate Synthase Inhibitors: Antibacterial Drug Leads. J. Med. Chem. 2014, 57 (13), 5693–5701. 10.1021/jm5004649.

(12) Ohlson, S.; Duong-Thi, M.-D. Fragment Screening for Drug Leads by Weak Affinity Chromatography (WAC-MS). Methods 2018, 146, 26–38. 10.1016/j.ymeth.2018.01.011.

(13) Lange, R.; Locher, H.; Wyss, P.; Then, R. The Targets of Currently Used Antibacterial Agents: Lessons for Drug Discovery. CPD 2007, 13 (30), 3140–3154. 10.2174/138161207782110408.

(14) Lee, L. V.; Granda, B.; Dean, K.; Tao, J.; Liu, E.; Zhang, R.; Peukert, S.; Wattanasin, S.; Xie, X.; Ryder, N. S.; Tommasi, R.; Deng, G. Biophysical Investigation of the Mode of Inhibition of Tetramic Acids, the Allosteric Inhibitors of Undecaprenyl Pyrophosphate Synthase. Biochemistry 2010, 49 (25), 5366– 5376. 10.1021/bi100523c.

(15) Jhoti, H.; Williams, G.; Rees, D. C.; Murray, C. W. The “rule of Three” for Fragment-Based Drug Discovery: Where Are We Now? Nat Rev Drug Discov 2013, 12 (8), 644–644. 10.1038/nrd3926-c1.

(16) Collins, P. M.; Ng, J. T.; Talon, R.; Nekrosiute, K.; Krojer, T.; Douangamath, A.; Brandao-Neto, J.; Wright, N.; Pearce, N. M.; Von Delft, F. Gentle, Fast and Effective Crystal Soaking by Acoustic Dispensing. Acta Crystallogr D Struct Biol 2017, 73 (3), 246–255. 10.1107/S205979831700331X.

(17) Hassell, A. M.; An, G.; Bledsoe, R. K.; Bynum, J. M.; Carter, H. L.; Deng, S.-J. J.; Gampe, R. T.; Grisard, T. E.; Madauss, K. P.; Nolte, R. T.; Rocque, W. J.; Wang, L.; Weaver, K. L.; Williams, S. P.; Wisely, G. B.; Xu, R.; Shewchuk, L. M. Crystallization of Protein–Ligand Complexes. Acta Crystallogr D Biol Crystallogr 2007, 63 (1), 72–79. 10.1107/S0907444906047020.

(18) Jukič, M.; Rožman, K.; Sova, M.; Barreteau, H.; Gobec, S. Anthranilic Acid Inhibitors of Undecaprenyl Pyrophosphate Synthase (UppS), an Essential Enzyme for Bacterial Cell Wall Biosynthesis. Front. Microbiol. 2019, 9, 3322. 10.3389/fmicb.2018.03322.

(19) Fujihashi, M.; Zhang, Y.-W.; Higuchi, Y.; Li, X.-Y.; Koyama, T.; Miki, K. Crystal Structure of *Cis* -Prenyl Chain Elongating Enzyme, Undecaprenyl Diphosphate Synthase. Proc. Natl. Acad. Sci. U.S.A. 2001, 98 (8), 4337–4342. 10.1073/pnas.071514398.

(20) Sinko, W.; De Oliveira, C.; Williams, S.; Van Wynsberghe, A.; Durrant, J. D.; Cao, R.; Oldfield, E.; McCammon, J. A. Applying Molecular Dynamics Simulations to Identify Rarely Sampled Ligand-bound Conformational States of Undecaprenyl Pyrophosphate Synthase, an Antibacterial Target. Chem Biol Drug Des 2011, 77 (6), 412–420. 10.1111/j.1747-0285.2011.01101.x.

(21) Ko, T.-P.; Chen, Y.-K.; Robinson, H.; Tsai, P.-C.; Gao, Y.-G.; Chen, A. P.-C.; Wang, A. H.-J.; Liang, P.-H. Mechanism of Product Chain Length Determination and the Role of a Flexible Loop in Escherichia Coli Undecaprenyl-Pyrophosphate Synthase Catalysis. Journal of Biological Chemistry 2001, 276 (50), 47474–47482. 10.1074/jbc.M106747200.

(22) Wang, W.; Dong, C.; McNeil, M.; Kaur, D.; Mahapatra, S.; Crick, D. C.; Naismith, J. H. The Structural Basis of Chain Length Control in Rv1086. Journal of Molecular Biology 2008, 381 (1), 129–140. 10.1016/j.jmb.2008.05.060.

(23) Kuo, C.-J.; Guo, R.-T.; Lu, I.-L.; Liu, H.-G.; Wu, S.-Y.; Ko, T.-P.; Wang, A. H.- J.; Liang, P.-H. Structure-Based Inhibitors Exhibit Differential Activities against *Helicobacter Pylori* and *Escherichia Coli* Undecaprenyl Pyrophosphate Synthases. BioMed Research International 2008, 2008 (1), 841312. 10.1155/2008/841312.

(24) Liebschner, D.; Afonine, P. V.; Moriarty, N. W.; Poon, B. K.; Sobolev, O. V.; Terwilliger, T. C.; Adams, P. D. Polder Maps: Improving OMIT Maps by Excluding Bulk Solvent. Acta Crystallogr D Struct Biol 2017, 73 (2), 148–157. 10.1107/S2059798316018210.

(25) Guo, R.-T.; Ko, T.-P.; Chen, A. P.-C.; Kuo, C.-J.; Wang, A. H.-J.; Liang, P.-H. Crystal Structures of Undecaprenyl Pyrophosphate Synthase in Complex with Magnesium, Isopentenyl Pyrophosphate, and Farnesyl Thiopyrophosphate. Journal of Biological Chemistry 2005, 280 (21), 20762–20774. 10.1074/jbc.M502121200.

(26) Lahra, M. M.; Ryder, N.; Whiley, D. M. A New Multidrug-Resistant Strain of *Neisseria Gonorrhoeae* in Australia. N Engl J Med 2014, 371 (19), 1850–1851. 10.1056/NEJMc1408109.

(27) Kabsch, W. *XDS*. Acta Crystallogr D Biol Crystallogr 2010, 66 (2), 125–132. 10.1107/S0907444909047337.

(28) Winn, M. D.; Ballard, C. C.; Cowtan, K. D.; Dodson, E. J.; Emsley, P.; Evans, P. R.; Keegan, R. M.; Krissinel, E. B.; Leslie, A. G. W.; McCoy, A.; McNicholas, S. J.; Murshudov, G. N.; Pannu, N. S.; Potterton, E. A.; Powell, H. R.; Read, R. J.; Vagin, A.; Wilson, K. S. Overview of the *CCP* 4 Suite and Current Developments. Acta Crystallogr D Biol Crystallogr 2011, 67 (4), 235–242. 10.1107/S0907444910045749.

(29) Emsley, P.; Lohkamp, B.; Scott, W. G.; Cowtan, K. Features and Development of *Coot*. Acta Crystallogr D Biol Crystallogr 2010, 66 (4), 486–501. 10.1107/S0907444910007493.

