## Supplementary Figures S1-S8 and Supplementary Tables S1 and S2 (PDF) for "Fragment Screening by Weak Affinity Chromatography Identifies Ligands for *Neisseria gonorrhoeae* Undecaprenyl Diphosphate Synthase"

[1] - Center of Medicinal Chemistry (CQMED), Center for Molecular Biology and Genetic Engineering (CBMEG), Universidade Estadual de Campinas, UNICAMP, 13083-886-Campinas, SP, Brazil.
[2] - Federal Institute of Education, Science and Technology of São Paulo, IFSP, 05110-000-São Paulo, SP, Brazil.
[3] - Eurofarma Laboratórios S/A, 06696-000-Itapevi, SP, Brazil.
[4] - Department of Organic Chemistry, Institute of Chemistry, Universidade Estadual de Campinas, UNICAMP, 13083-970-Campinas, SP, Brazil
[5] - Structural Genomics Consortium and Division of Chemical Biology and Medicinal Chemistry, UNC Eshelman School of Pharmacy, University of North Carolina, Chapel Hill, North Carolina 27599, USA.

### These authors contributed equally to this work.

#### Supplementary Figure S1

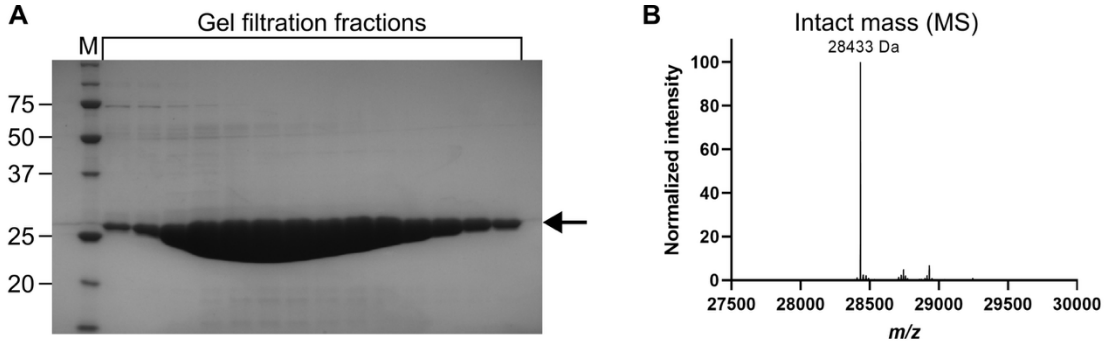

**Figure S1.** Purification of recombinant *Ng*UPPS. (A) SDS-PAGE analysis of fractions from the final purification step (size-exclusion chromatography). The arrow marks the expected molecular weight of *Ng*UPPS. M, molecular weight marker (band sizes indicated in kDa). (B) Intact mass spectrometry confirming the expected molecular weight of purified *Ng*UPPS (28,432 Da).

#### Supplementary Figure S2

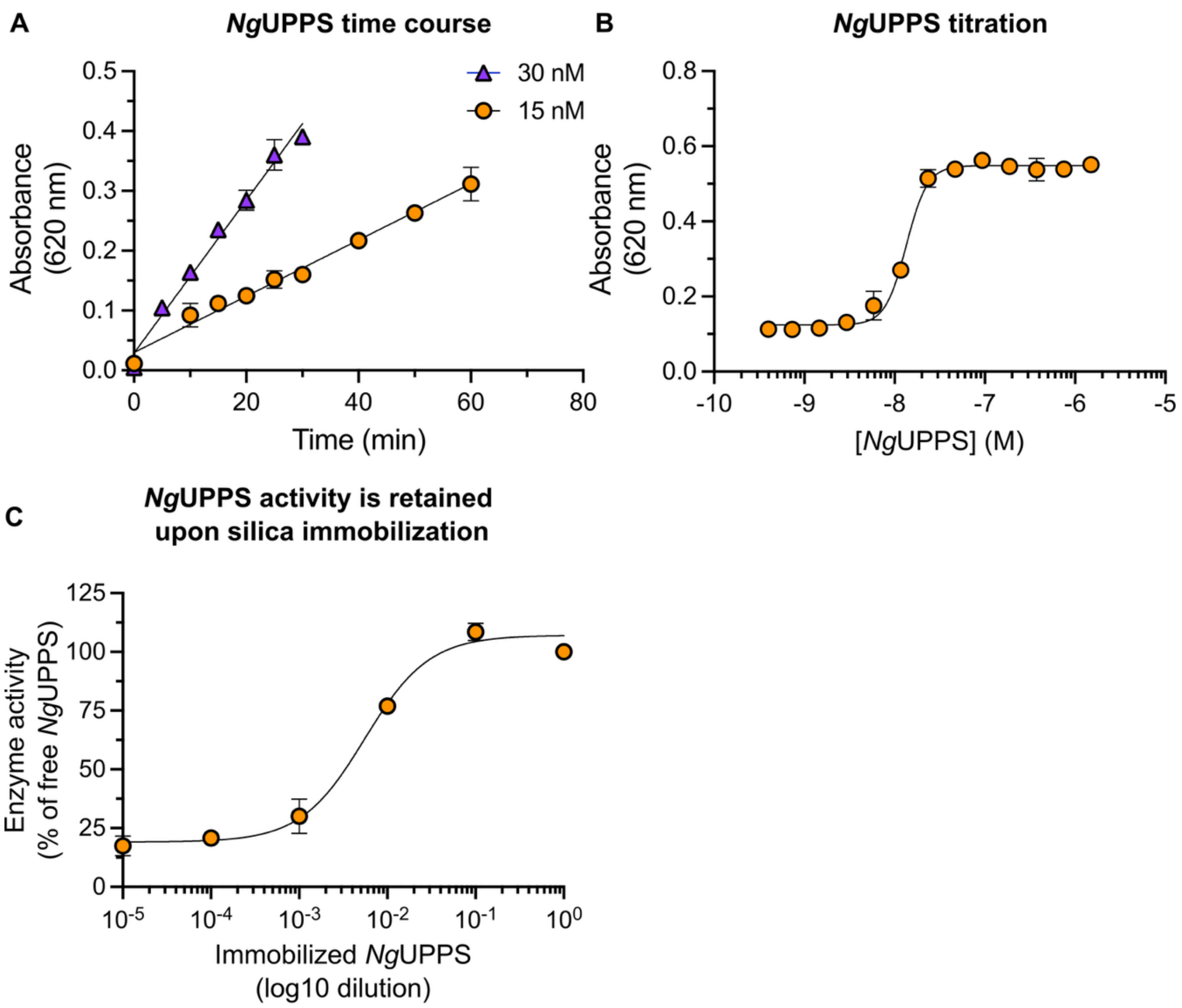

**Figure S2.** Enzymatic assay development for *Ng*UPPS. (A) Time course of product formation using 15 nM or 30 nM purified *Ng*UPPS. Data represent the mean ± SEM from a representative experiment performed in duplicate. The lines indicate linear fits to the experimental data, with $R^2$ values of 0.97 for 15 nM and 0.98 for 30 nM. (B) Enzyme titration showing *Ng*UPPS activity across a range of concentrations. Data represent the mean ± SEM from a representative experiment performed in duplicate. The line represents a four-parameter sigmoidal fit to the data. The calculated $IC_{80}$ value is 20 nM. (C) Enzymatic activity of silica-immobilized *Ng*UPPS across a range of dilutions, normalized to the activity of the free enzyme at 10 nM (100%) and silica alone (0%). Data represent mean ± SEM from a representative experiment performed in duplicate. The line represents a four-parameter sigmoidal fit to the data.

#### Supplementary Figure S3

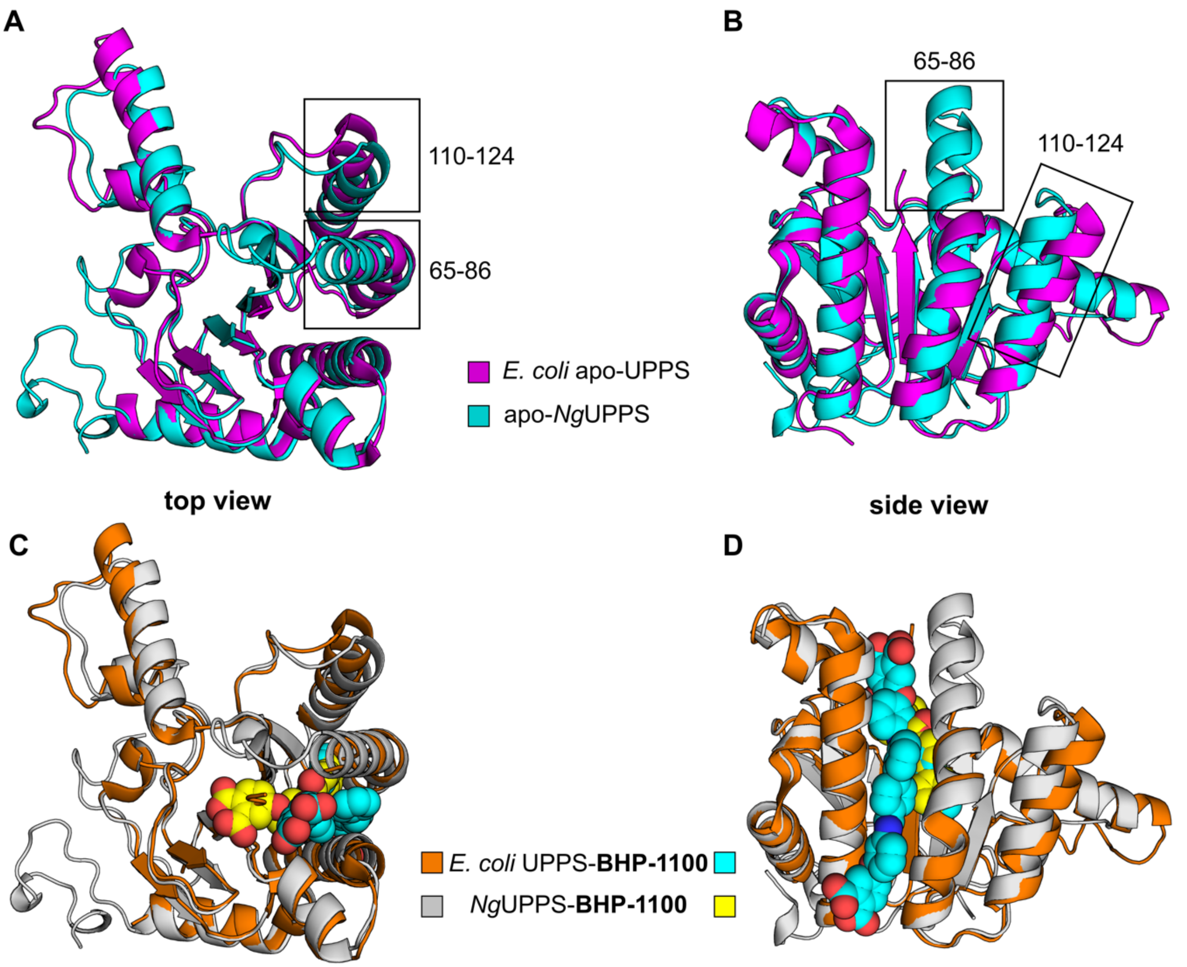

**Figure S3.** Comparison of apo- and **BHP-1100**-bound *E. coli* and *N. gonorrhoeae* UPPS crystal structures. (A, B) Cartoon representations of the apo *E. coli* structure (magenta; PDB ID: 3QAS) and the apo-*Ng*UPPS structure (cyan; PDB ID: 9OH8). Boxes highlight regions in *E. coli* UPPS that are either disordered or structurally divergent relative to the corresponding regions in *Ng*UPPS. Residue numbers correspond to the *Ng*UPPS sequence. (C, D) Cartoon representations of the **BPH-1100**–bound co-structures of *E. coli* (orange; PDB ID: 3SGX) and *Ng*UPPS (gray; PDB ID: 9OHA). **BPH-1100** is shown as spheres, with ligand carbon atoms colored yellow for *Ng*UPPS and cyan for *E. coli* UPPS. Structures superimposed using PyMol.

#### Supplementary Figure S4

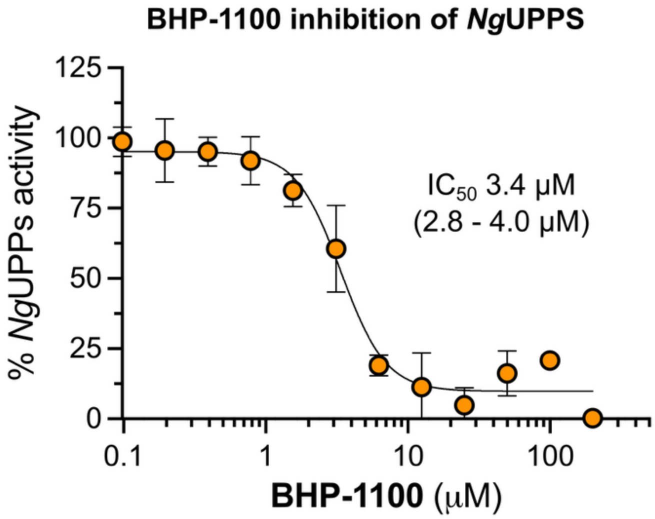

**Figure S4.** Dose–response curve for **BHP-1100** inhibition of *Ng*UPPS. Data represent the mean ± standard error of the mean (SEM) from two independent experiments performed in duplicate. The $IC_{50}$ value was determined by fitting the data to a four-parameter sigmoidal dose–response model; 95% confidence intervals are shown in parentheses.

#### Supplementary Figure S5

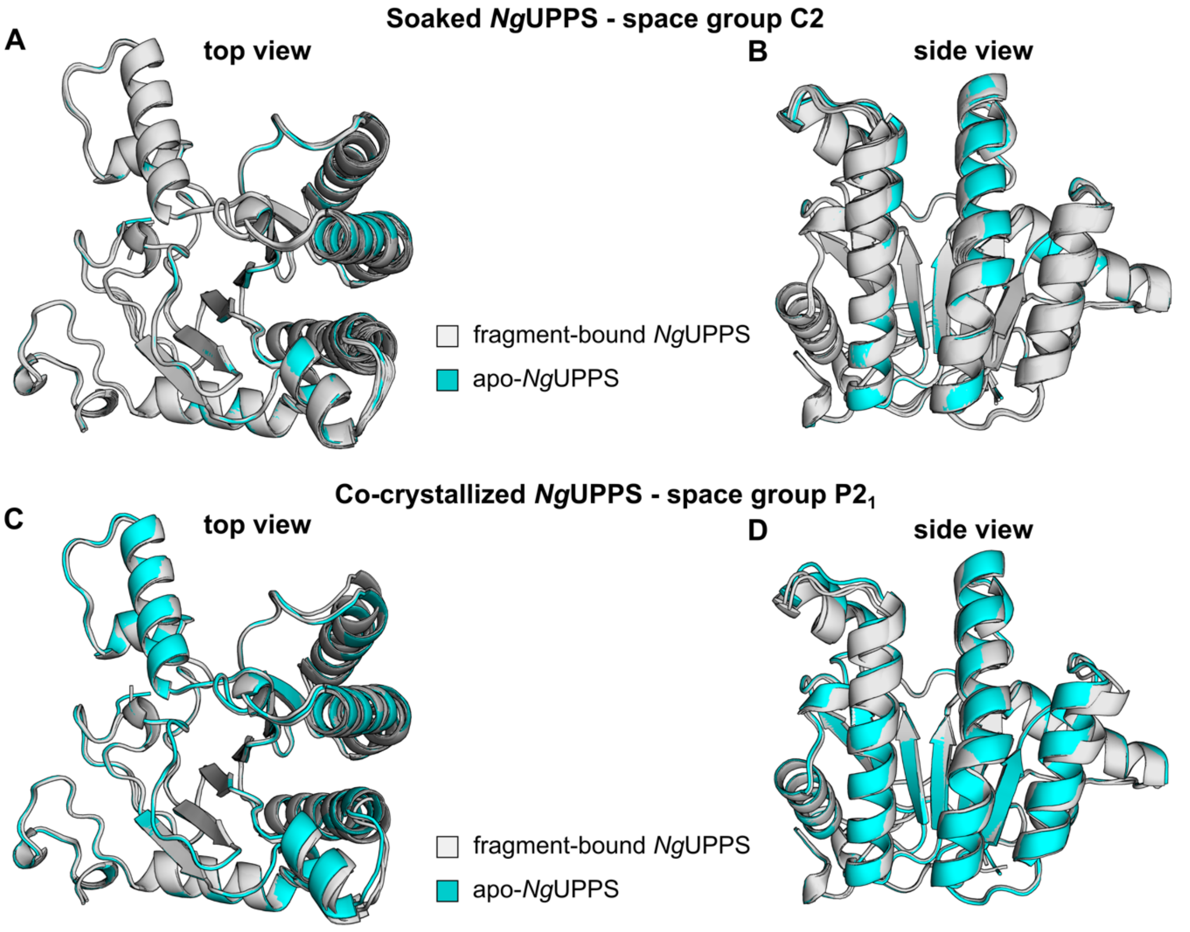

**Figure S5**. Comparison of apo- and fragment-bound *Ng*UPPS crystal structures. (A, B) Cartoon representations of apo-*Ng*UPPS (cyan) overlaid with the 15 fragment-bound *Ng*UPPS structures crystallized in the C2 space group (gray). (C, D) Cartoon representations of apo-*Ng*UPPS (cyan) overlaid with the two protomers from the fragment-bound *Ng*UPPS structure crystallized in the P2$_1$ space group (gray). Structures superimposed using PyMol. PDB IDs of apo-*Ng*UPPS and *Ng*UPPS bound to fragments are: 9OH8 (Apo) 9OI4 (**1R-0197**), 9OHI (**PS-5549**), 9OHJ (**4R-0623**), 9OIW (**GF-0701**), 9OHO (**PS-3756**), 9OHB (**PS-5485**) and 9OHG (**PS-5570**), 9OJE (**5Br-P**), 9OI6 (**FD-0739**), 9OIY (**DA-0841**), 9OIV (**4W-0801**), 9OJB (**PS-0394**), 9OI5 (**PS-4845**), 9OHP (**BA-0843**), 9OJ1 (**DA-0854**), 9OJA (**FC-0609**).

#### Supplementary Figure S6

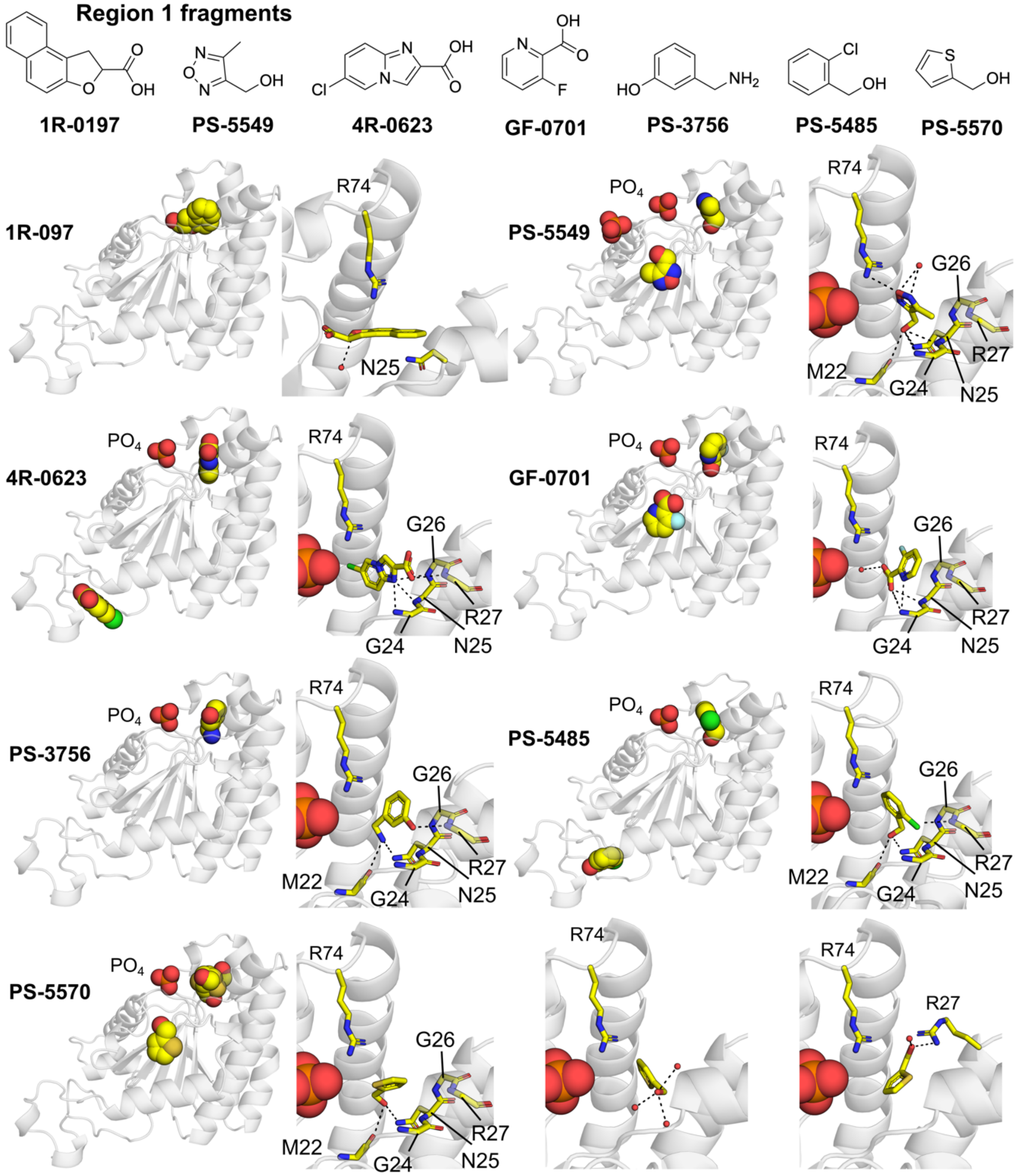

**Figure S6**. Structure and binding details of fragments bound to *Ng*UPPS Region 1. The protein is shown as a gray, semi-transparent cartoon. Fragments and phosphate ions are depicted as spheres in the zoomed-out panels; fragments bound in Region 1 are shown as sticks in the zoomed-in views. Black dashed lines indicate potential hydrogen bonds. Fragments that also occupy a secondary site are shown only in the zoomed-out panels. PDB IDs of *Ng*UPPS bound to fragments are: 9OI4 (**1R-0197**), 9OHI (**PS-5549**), 9OHJ (**4R-0623**), 9OIW (**GF-0701**), 9OHO (**PS-3756**), 9OHB (**PS-5485**) and 9OHG (**PS-5570**).

#### Supplementary Figure S7

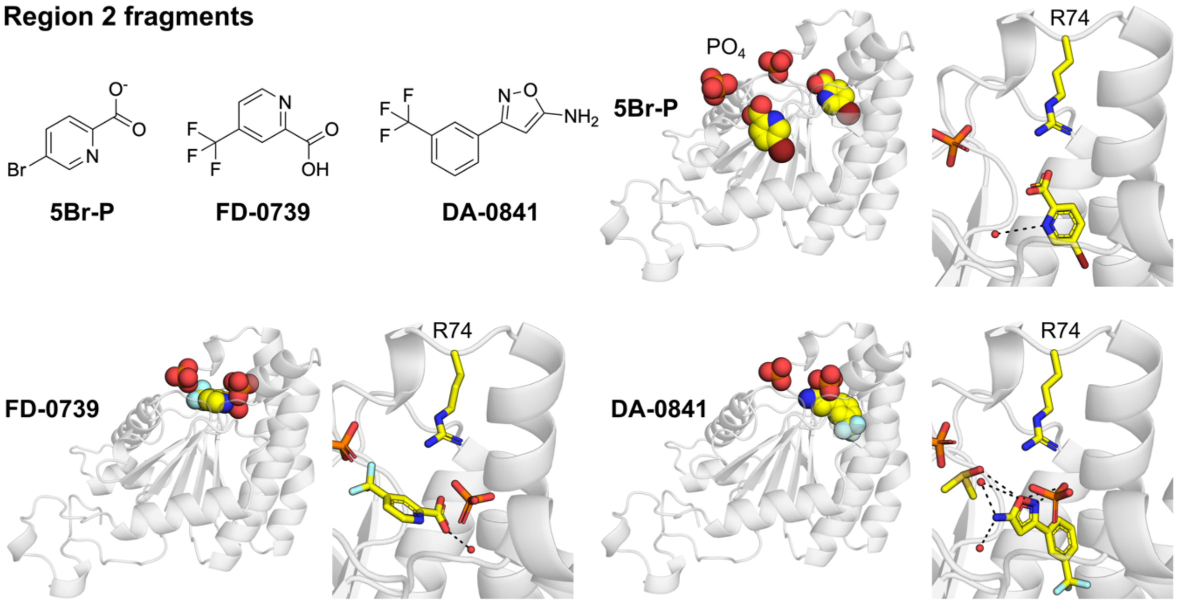

**Figure S7**. Structure and binding details of fragments bound to *Ng*UPPS Region 2. The protein is shown as a gray, semi-transparent cartoon. Fragments and phosphate ions are depicted as spheres in the zoomed-out panels; fragments bound in Region 2 are shown as sticks in the zoomed-in views. Black dashed lines indicate potential hydrogen bonds. Fragments that also occupy a secondary site are shown only in the zoomed-out panels. PDB IDs of *Ng*UPPS bound to fragments are: 9OJE (**5Br-P**), 9OI6 (**FD-0739**), 9OIY (**DA-0841**).

#### Supplementary Figure S8

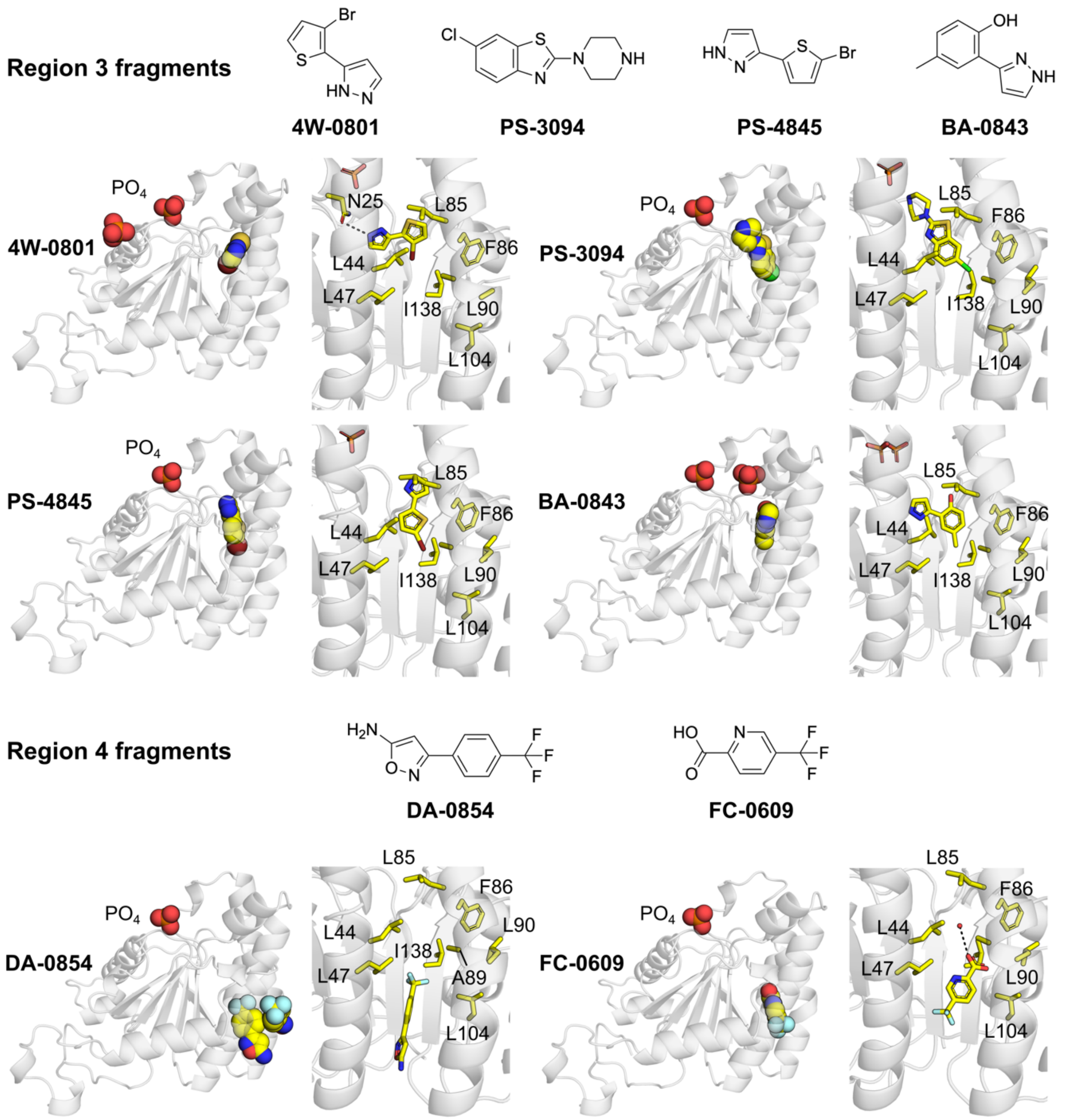

**Figure S8**. Structure and binding details of fragments bound to *Ng*UPPS Regions 3 and 4. The protein is shown as a gray, semi-transparent cartoon. Fragments and phosphate ions are depicted as spheres in the zoomed-out panels; fragments bound in Regions 3 and 4 are shown as sticks in the zoomed-in views. Black dashed lines indicate potential hydrogen bonds. Fragments that also occupy a secondary site are shown only in the zoomed-out panels. PDB IDs of *Ng*UPPS bound to fragments are: 9OIV (**4W-0801**), 9OJB (**PS-3094**), 9OI5 (**PS-4845**), 9OHP (**BA-0843**), 9OJ1 (**DA-0854**), 9OJA (**FC-0609**).

**Supplementary Table S1**

| Table S1 – Specific retention times ($t_{spec}$, min) of selected fragments | | | | | |
|---|---|---|---|---|---|
| Fragment ID | SMILES | $t_{spec}$ (min) | | | |
| | | n=1 | n=2 | Avg[a] | StdDev[a] |
| PS-5485 | OCC1=C(Cl)C=CC=C1 | 18.45 | 18.45 | 18.45 | 0.00 |
| PS-5570 | OCC1=CC=CS1 | 18.40 | 18.40 | 18.40 | 0.00 |
| PS-5549 | CC1=NON=C1CO | 18.35 | 18.35 | 18.35 | 0.00 |
| 1R-0197 | OC(C1CC2=C3C=CC=CC3=CC=C2O1)=O | 18.10 | 18.10 | 18.10 | 0.00 |
| PS-4305 | OC(C1=CC=NN1C2=CC=CC=C2)=O | 16.10 | 16.10 | 16.10 | 0.00 |
| FD-0739 | OC(C1=NC=CC(C(F)(F)F)=C1)=O | 16.05 | 16.05 | 16.05 | 0.00 |
| BA-0843 | CC1=CC(C2=NNC=C2)=C(O)C=C1 | 11.86 | 11.86 | 11.86 | 0.00 |
| 5Br-P | OC(C1=NC=C(C=C1)Br)=O | 10.30 | 10.30 | 10.30 | 0.00 |
| 4R-0623 | OC(C1=CN2C=C(C=CC2=N1)Cl)=O | 9.40 | 9.40 | 9.40 | 0.00 |
| PS-3756 | NCC1=CC(O)=CC=C1 | 9.05 | 9.00 | 9.03 | 0.04 |
| PS-4845 | BrC1=CC=C(C2=NNC=C2)S1 | 8.70 | 8.70 | 8.70 | 0.00 |
| PS-5844 | CN(CCN1CCNCC1)C | 8.00 | 9.00 | 8.50 | 0.71 |
| GF-0701 | OC(C1=NC=CC=C1F)=O | 7.80 | 7.70 | 7.75 | 0.07 |
| DA-0854 | NC1=CC(C2=CC=C(C(F)(F)F)C=C2)=NO1 | 6.65 | 6.65 | 6.65 | 0.00 |
| 4W-0802 | C12=C(N3C=CC=C3)NN=C1C=CC=C2 | 6.30 | 6.30 | 6.30 | 0.00 |
| FC-0609 | OC(C1=NC=C(C(F)(F)F)C=C1)=O | 6.25 | 6.25 | 6.25 | 0.00 |
| PS-3094 | ClC1=CC2=C(N=C(N3CCNCC3)S2)C=C1 | 5.70 | 5.90 | 5.80 | 0.14 |
| PS-5127 | CC1=CC2=C(NN=C2)C(Br)=C1 | 5.35 | 5.35 | 5.35 | 0.00 |
| MI-0621 | CN1C=C(C(O)=O)N=C1 | 5.30 | 5.30 | 5.30 | 0.00 |
| 3W-0219 | OC1=CN(C2=CC(C(F)(F)F)=CC=C2)N=C1 | 5.15 | 5.15 | 5.15 | 0.00 |
| FS-1153 | FC(OC1=C(CC#N)C=CC=C1)F | 5.10 | 5.10 | 5.10 | 0.00 |
| DA-0841 | NC1=CC(C2=CC(C(F)(F)F)=CC=C2)=NO1 | 5.10 | 5.10 | 5.10 | 0.00 |
| 4W-0801 | BrC1=C(C2=NNC=C2)SC=C1 | 5.10 | 5.10 | 5.10 | 0.00 |

[a] average (Avg.) and standard deviations (StdDev) shown for two independent experiments.

**Table S2**

| | PS-5485 | PS-5570 | PS-5549 |
|---|---|---|---|
| **PDB code** | **9OHB** | **9OHG** | **9OHI** |
| **Ligand Entry (RegID)** | | | |
| **Data collection** | | | |
| Space group | C2 | C2 | C2 |
| Cell dimensions | | | |
| *a*, *b*, *c* (Å) | *115.58 50.47 47.28* | *115.420 50.547 47.014* | *115.035 50.117 47.150* |
| α, β, γ (°) | *90.00 106.97 90.00* | *90 106.76 90.00* | *90.00 107.09 90.00* |
| Resolution (Å) | 45.91-1.53 (8.22-1.5) | 45.97-1.57 (8.60-1.57) | 45.60 (1.79) |
| $R_{sym}$ or $R_{merge}$ | 0.044(0.918) | 0.048(0.647) | 0.064(0.877) |
| CC1/2 | 0.99(0.756) | 0.999(0.810) | 0.999(0.794) |
| I / σI | 18.1(2) | 17.7(2.6) | 6.6(6.9) |
| Completeness (%) | 99.1(97.9) | 98.8(97.7) | 100.0(100.0) |
| Redundancy | 6.7(6.8) | 6.7(6.6) | 6.6(6.9) |
| **Refinement** | | | |
| Resolution (Å) | 55.27-1.50 | 46.01(1.57) | 45.65(1.79) |
| No. reflections | 59404 | 34049 | 23123 |
| $R_{work}$ / $R_{free}$(%) | 0.1912/0.2154 | 0.17398/0.20197 | 0.17758/0.22026 |
| No. atoms | | | |
| Protein | 3863 | 3867 | 3799 |
| Ligand/ion | 16 | 13 | 14 |
| Water | 128 | 169 | 149 |
| *B*-factors | | | |
| Protein | 26.774 | 20.6 | 31.55 |
| Ligand/ion | 49.76 | 33.41 | 50.62 |
| Water | 37.96 | 38.58 | 43.11 |
| R.m.s. deviations | | | |
| Bond lengths (Å) | 0.01 | 0.0103 | 0.008 |
| Bond angles (°) | 1.583 | 1.922 | 1.428 |

*Values in parentheses are for highest-resolution shell.

| | 4R-0623 | PS-3756 | PS-4845 |
|---|---|---|---|
| **PDB code** | **9OHJ** | **9OHO** | **9OI5** |
| **Ligand Entry (RegID)** | | | |
| **Data collection** | | | |
| Space group | C2 | C2 | C2 |
| Cell dimensions | | | |
| *a*, *b*, *c* (Å) | 115.36 50.57 47.17 | 116.11 50.34 47.35 | 115.203 50.115 47.291 |
| α, β, γ (°) | 90.00 107.06 90.00 | 90.00 106.75 90.00 | 90.000 107.132 90.000 |
| Resolution (Å) | 45.97-1.90 (9.11 1.86) | 45.86-1.58 (8.49-1.55) | 45.61 1.84 ( 9.00-1.80) |
| $R_{sym}$ or $R_{merge}$ | 0.086 (0.890) | 0.046 (0.856) | 0.064 (0.818) |
| CC1/2 | 0.999 (0.816) | 0.999 (0.750) | 0.999 (0.800) |
| I / σI | 12.7 (2.4) | 18.1 (2.0) | 15.2 (2.3) |
| Completeness (%) | 99.8 (99.6) | 99.9 (100.0) | 99.6 (98.9) |
| Redundancy | 6.6 (6.9) | 6.6-6.5 | 6.6 (6.9) |
| **Refinement** | | | |
| Resolution (Å) | 46.1-1.9 (9.11-1.86) | 45.90-1.55 | 45.6-1.8 |
| No. reflections | 20864 | 36201 | 26132 |
| $R_{work}$ / $R_{free}$(%) | 0.17122/0.21596 | 0.1771/0.2017 | 0.1783/0.2258 |
| No. atoms | | | |
| Protein | 3834 | 3858 | 3819 |
| Ligand/ion | 18 | 18 | 16 |
| Water | 95 | 140 | 104 |
| *B*-factors | | | |
| Protein | 33.8 | 28.63 | 32.21 |
| Ligand/ion | 70.13 | 52.20 | 113.74 |
| Water | 41.53 | 38.27 | 39.50 |
| R.m.s. deviations | | | |
| Bond lengths (Å) | 0.0075 | 0.0109 | 0.0076 |
| Bond angles (°) | 1.591 | 1.794 | 1.628 |

*Values in parentheses are for highest-resolution shell.

| | PS-5844 | GF-0701 | DA-0854 |
|---|---|---|---|
| **PDB code** | **9OIE** | **9OIW** | **9OJ1** |
| **Ligand Entry (RegID)** | | | |
| **Data collection** | | | |
| Space group | C2 | C2 | C2 |
| Cell dimensions | | | |
| *a*, *b*, *c* (Å) | 115.31 50.10 47.16 | 115.18 50.46 47.08 | 114.96 50.46 47.11 |
| α, β, γ (°) | 90.00 107.06 90.00 | 90.00 106.61 90.00 | 90.00 106.69 90.00 |
| Resolution (Å) | 45.61-1.84 ( 9.00-1.80) | 45.89-1.78 ( 9.09-1.75) | 45.88-1.58 (8.49-1.55) |
| $R_{sym}$ or $R_{merge}$ | 0.075 (0.683) | 0.061 (0.794) | 0.050 (0.820) |
| CC1/2 | 0.998 (0.864) | 0.999 (0.764) | 0.999 (0.797) |
| I / σI | 12.7 (2.4) | 15.9 (2.3) | 17.4 (2.2) |
| Completeness (%) | 99.8 (100) | 99.9 (100) | 96.9 (95.2) |
| Redundancy | 6.4 (6.9) | 6.6 (7.0) | 6.8 (6.9) |
| **Refinement** | | | |
| Resolution (Å) | 55.12-1.80 | 45.94-1.75 | 45.92-1.55 |
| No. reflections | 22756 | 24878 | 34607 |
| $R_{work}$ / $R_{free}$(%) | 0.1827/ 0.2260 | 0.17827 / 0.21827 | 0.17403 / 0.19832 |
| No. atoms | | | |
| Protein | 3834 | 3811 | 3875 |
| Ligand/ion | 30 | 14 | 23 |
| Water | 100 | 127 | 132 |
| *B*-factors | | | |
| Protein | 31.5 | 30.03 | 27.47 |
| Ligand/ion | 55.90 | 58.202 | 81.60 |
| Water | 30.12 | 40.84 | 36.38 |
| R.m.s. deviations | | | |
| Bond lengths (Å) | 0.0083 | 0.009 | 0.01 |
| Bond angles (°) | 1.643 | 1.686 | 1.773 |

*Values in parentheses are for highest-resolution shell.

| | FC-0609 | PS-3094 | DA-0841 |
|---|---|---|---|
| **PDB code** | **9OJA** | **9OJB** | **9OIY** |
| **Ligand Entry (RegID)** | | | |
| **Data collection** | | | |
| Space group | C2 | C2 | C2 |
| Cell dimensions | | | |
| *a*, *b*, *c* (Å) | 115.43 50.24 47.40 | 116.132 50.397 47.383 | 116.15 50.36 47.37 |
| α, β, γ (°) | 90.00 106.89 90.00 | 90.00 106.90 90.00 | 90.00 106.77 90.00 |
| Resolution (Å) | 45.73-1.42 (7.67-1.40) | 45.900-1.47 (7.94-1.45) | 45.88-1.51 ( 8.11 1.48) |
| $R_{sym}$ or $R_{merge}$ | 0.045 (1.003) | 0.074 (1.036) | 0.039 (0.656) |
| CC1/2 | 0.999 (0.669) | 0.998 (0.804) | 0.999 (0.846) |
| I / σI | 18.2 (1.8) | 13.8 (2.1) | 20.0 (2.7) |
| Completeness (%) | 98.5 (92.7) | 99.0 (96.5) | 100 (100) |
| Redundancy | 6.7 (6.8) | 6.7 (6.8) | 6.6 (6.7) |
| **Refinement** | | | |
| Resolution (Å) | 55.23-1.4 | 45.94-1.45 | 55.61-1.48 |
| No. reflections | 71114 | 43728 | 41578 |
| $R_{work}$ / $R_{free}$(%) | 0.1735 / 0.1947 | 0.19694 / 0.22747 | 0.1815 / 0.2100 |
| No. atoms | | | |
| Protein | 3859 | 3832 | 1924 |
| Ligand/ion | 17 | 28 | 23 |
| Water | 180 | 193 | 187 |
| *B*-factors | | | |
| Protein | 23.35 | 22.11 | 25.05 |
| Ligand/ion | 72.73 | 49.5 | 70.97 |
| Water | 34.29 | 33.05 | 38.20 |
| R.m.s. deviations | | | |
| Bond lengths (Å) | 0.0118 | 0.01 | 0.0104 |
| Bond angles (°) | 1.832 | 1.878 | 1.763 |

*Values in parentheses are for highest-resolution shell.

| | FD-0739 | 1R-0197 | BA-0843 |
|---|---|---|---|
| **PDB code** | **9OI6** | **9OI4** | **9OHP** |
| **Ligand Entry (RegID)** | | | |
| **Data collection** | | | |
| Space group | C2 | P 1 21 1 | C2 |
| Cell dimensions | | | |
| *a*, *b*, *c* (Å) | 116.071 50.193 47.288 | 47.340 57.459 100.024 | 115.669 50.006 47.216 |
| α, β, γ (°) | 90.00 106.84 90.00 | 90.00 96.61 90.00 | 90.00 107.14 90.00 |
| Resolution (Å) | 45.74-2.38 (8.9-2.30) | 47.02-2.37 (8.53 2.28) | 45.56-1.61 (8.65-1.58) |
| $R_{sym}$ or $R_{merge}$ | 0.130 (0.676) | 0.193 (0.921) | 0.044 (0.842) |
| CC1/2 | 0.996 (0.785) | 0.990 (0.732) | 0.999 (0.757) |
| I / σI | 8.1 (2.2) | 7.0 (1.9) | 19.2 (2.2) |
| Completeness (%) | 99.9 (100) | 100 (100) | 100 (100) |
| Redundancy | 5.5 (5.3) | 6.6 (7.0) | 6.6 (6.7) |
| **Refinement** | | | |
| Resolution (Å) | 45.78 (2.3) | 47.07 (2.28) | 45.60 (1.58) |
| No. reflections | 11167 | 23358 | 33606 |
| $R_{work}$ / $R_{free}$(%) | 0.20141 / 0.27185 | 0.18404 / 0.24561 | 0.17798 / 0.19811 |
| No. atoms | | | |
| Protein | 1929 | 3941 | 1953 |
| Ligand/ion | 17 | 26 | 23 |
| Water | 46 | 161 | 146 |
| *B*-factors | | | |
| Protein | 43.56 | 31.49 | 28.10 |
| Ligand/ion | 97.7 | 47.29 | 43.92 |
| Water | 41.91 | 29.04 | 36.54 |
| R.m.s. deviations | | | |
| Bond lengths (Å) | 0.0068 | 0.0083 | 0.01 |
| Bond angles (°) | 1.626 | 1.807 | 1.716 |

*Values in parentheses are for highest-resolution shell.

| | 5-BrPIc | 4W-0801 | Apo protein |
|---|---|---|---|
| **PDB code** | **9OJE** | **9OIV** | **9OH8** |
| **Ligand Entry (RegID)** | | | |
| **Data collection** | | | |
| Space group | C2 | C2 | C2 |
| Cell dimensions | | | |
| *a*, *b*, *c* (Å) | 115.315 50.263 47.245 | 116.15 49.77 46.78 | 116.76 50.20 47.65 |
| α, β, γ (°) | 90.00 106.86 90.00 | 90.00 106.35 90.00 | 90.00 106.85 90.00 |
| Resolution (Å) | 45.74-1.82 (9.08-1.78) | 45.44-2.05 (8.94-2.00) | 19.55-1.68 (9.04-1.65) |
| $R_{sym}$ or $R_{merge}$ | 0.070 (0.708) | 0.063(0.700) | 0.063 ( 1.180) |
| CC1/2 | 0.999 (0.815) | 0.999 (0.886) | 0.999 (0.728) |
| I / σI | 15.2 (2.7) | 15.1 (2.4) | 14.8 (1.6) |
| Completeness (%) | 99.9 (99.9) | 99.9 (100) | 99.9 (100) |
| Redundancy | 6.6 (6.9) | 6.5 (6.6) | 6.7 (7.0) |
| **Refinement** | | | |
| Resolution (Å) | 45.78 (1.78) | 45.48 (2.0) | 20.0 (1.65) |
| No. reflections | 23690 | 16591 | 44680 |
| $R_{work}$ / $R_{free}$(%) | 0.16650 / 0.20507 | 0.19477 / 0.25694 | 0.16535 / 0.1900 |
| No. atoms | | | |
| Protein | 3784 | 1936 | 1918 |
| Ligand/ion | 14 | 16 | N/A |
| Water | 172 | 170 | 325 |
| *B*-factors | | | |
| Protein | 27.45 | 44.197 | 34.894 |
| Ligand/ion | 64.6 | 66.69 | N/A |
| Water | 30.40 | 50.26 | 39.10 |
| R.m.s. deviations | | | |
| Bond lengths (Å) | 0.009 | 0.008 | 0.01 |
| Bond angles (°) | 1.646 | 1.85 | 1.704 |

*Values in parentheses are for highest-resolution shell.

| | BPH-1100 |
|---|---|
| **PDB code** | **9OHA** |
| **Ligand Entry (RegID)** | |
| **Data collection** | |
| Space group | C2 |
| Cell dimensions | |
| *a*, *b*, *c* (Å) | 115.315 50.263 47.245 |
| α, β, γ (°) | 90.00 106.86 90.00 |
| Resolution (Å) | 45.74-1.82 (9.08-1.78) |
| $R_{sym}$ or $R_{merge}$ | 0.070 (0.708) |
| CC1/2 | 0.999 (0.815) |
| I / σI | 15.2 (2.7) |
| Completeness (%) | 99.9 (99.9) |
| Redundancy | 6.6 (6.9) |
| **Refinement** | |
| Resolution (Å) | 45.34 (1.75) |
| No. reflections | 25035 |
| $R_{work}$ / $R_{free}$(%) | 0.18447 / 0.21272 |
| No. atoms | |
| Protein | 3836 |
| Ligand/ion | 53 |
| Water | 100 |
| *B*-factors | |
| Protein | 35.12 |
| Ligand/ion | 64.04 |
| Water | 40.82 |
| R.m.s. deviations | |
| Bond lengths (Å) | 0.007 |
| Bond angles (°) | 1.639 |

*Values in parentheses are for highest-resolution shell.
